# Cell-type-specific regulatory variation shapes maize heterosis

**DOI:** 10.64898/2026.08.20.744945

**Authors:** Luguang Jiang, Fabio Gomez-Cano, Juncheng Luo, Mark A.A. Minow, Alexandre P. Marand

## Abstract

Heterosis, the superior performance of hybrids over their parents, has been widely exploited to enhance crop productivity, but its underlying regulatory mechanisms remain incompletely understood. In particular, how *cis*-regulatory elements (CREs) vary between parents and hybrids, and how this variation is organized across cell types, remains largely unresolved. Here we profiled chromatin accessibility in 131,890 nuclei derived from seedlings of three *Zea mays* (maize) inbred lines and their reciprocal hybrids, resolving 14 major cell types. Parental haplotype comparisons showed that most accessible chromatin regions (ACRs) were sequence-conserved and have similar binarized chromatin accessibility status. At cellular resolution, hybridization broadly attenuated chromatin accessibility cell-type specificity, weakening parental cell-type bias and yielding a more even chromatin accessibility profile across cellular contexts. Chromatin inheritance was strongly cell-type dependent, with non-additive inheritance preferentially concentrated at cell-type-specific ACRs. Although *cis* effects were most prevalent overall, ACRs with attenuated cell-type specificity in hybrids were enriched for non-additive inheritance and *trans* effects. Attenuated ACRs were more prone to transcription factor (TF) footprint gains than loci retaining high cell-type specificity, with DNA-BINDING WITH ONE FINGER (DOF) and VASCULAR PLANT ONE ZINC FINGER (VOZ) among the motif families most enriched in high-confidence footprint-gaining events. ACRs with footprint gains were preferentially linked to genes involved in development, hormone responses and growth, including DOF-family gains at *GOLDEN2-like* (*GLK2*), where both the motif family and the locus have been implicated in bundle-sheath development and C4 photosynthetic specialization. Together, our findings reveal that parental chromatin accessibility patterns are reconfigured across cell types in maize hybrids, with attenuation of cell-type specificity offering a potential cellular mechanism underlying heterosis.

## INTRODUCTION

The success of plant breeding depends on how genetic variation is combined and expressed in progeny. Crosses between genetically distinct individuals not only generate new allele combinations but can also lead to heterosis, or hybrid vigor, where F1 hybrids outperform their parental lines for key agronomic traits such as growth, yield and environmental adaptability ^1,2^. Heterosis has underpinned crop improvement for over a century and continues to support global food security ^2–4^. Although three classical genetic models, including dominance, overdominance and epistasis, provide conceptual explanations for hybrid vigor ^5–12^, the molecular regulation of heterosis formation remains enigmatic. In particular, how genetic variation between parental genomes is combined, coordinated and deployed to produce superior hybrid performance from a molecular perspective remains an open question.

A central link between genetic variation and phenotypic diversity lies in the regulation of gene expression. Genetic variation between parental genomes can influence gene expression through multiple regulatory layers, including chromatin accessibility ^13–17^, epigenetic modifications ^18–23^ and 3D genome architecture ^17,24–27^. Recent advances in high-throughput omics technologies have enabled systematic, genome-wide interrogation of the molecular basis of heterosis across multiple regulatory layers. Among them, chromatin accessibility constitutes a key regulatory layer by modulating the ability of TFs and co-regulatory complexes to engage regulatory DNA and shape gene expression programs ^28^. Genome-wide chromatin accessibility profiling ^29^ has increasingly been used across diverse plant species to compare hybrids with their parents and to investigate how changes in chromatin accessibility contribute to altered gene activity in hybrids ^13–16^. A recent maize study using MNase hypersensitivity sequencing (MH-seq) showed that chromatin accessibility in F1 hybrids is largely conserved and additively inherited, while a small subset of transgressively upregulated ACRs may contribute to heterosis ^16^. Likewise, maize assay for transposase-accessible chromatin with sequencing (ATAC-seq) found predominantly additive chromatin accessibility inheritance and revealed *trans*-regulatory modulation in F1 hybrids that coincided with strong heterosis ^15^.

However, existing analyses of chromatin accessibility in hybrids have relied on bulk tissues, in which signals from heterogeneous cell populations are averaged. Such measurements obscure cell-type-specific regulatory programs and limit mechanistic insight into how parental regulatory variation is integrated within individual cellular contexts. Given that many agronomic traits underlying heterosis arise from coordinated activities of distinct cell types, resolving regulatory inheritance at cellular resolution is essential for elucidating the molecular basis of hybrid vigor. Recent advances in single-cell chromatin accessibility profiling provide a powerful means to address these limitations. Single-cell ATAC-seq (scATAC-seq) has been recently applied in plants to resolve cell-type-specific regulatory landscapes and *cis*-regulatory element usage at high resolution ^30–45^. Applying such approaches to both parents and F1 hybrids offers the opportunity to directly interrogate how parental chromatin accessibility states combine in different cell types, thereby linking cell-type regulatory element inheritance to hybrid phenotypes.

Here we used single-cell combinatorial fluidic indexing ATAC-sequencing (scifi-ATAC-seq) ^46^ to profile chromatin accessibility across three maize inbred lines and their six reciprocal F1 hybrids at single-cell resolution. By integrating cell-type-resolved, haplotype-aware and allele-specific chromatin accessibility analyses, we show that hybridization broadly attenuates parental cell-type-specific chromatin accessibility through diffuse deployment of regulatory activity across hybrid cell types, rather than through changes in cell-type composition. Chromatin accessibility was predominantly constant or additively inherited, whereas non-additive inheritance and *trans*-involved regulatory effects were concentrated at highly cell-type-specific loci and at regulatory elements with attenuated chromatin accessibility cell-type specificity in hybrids. These loci also exhibited TF footprint gains near genes involved in growth development and hormone responses. Collectively, our study reveals how parental chromatin accessibility is integrated across cellular contexts and identifies attenuation of cell-type specificity as a pervasive feature of regulatory remodeling in maize hybrids, providing cell-type-resolved insights into the contribution of chromatin accessibility to heterosis.

## RESULTS

### Single-cell chromatin accessibility across maize inbreds and hybrids

To advance our understanding of the regulatory programs that promote maize heterosis, we performed scifi-ATAC-seq ^46^ on seven-day-old maize seedlings collected from three inbred lines (B73, Ki3, and Oh43) and their six reciprocal hybrids with two biological replicates (**Fig. 1a and Supplementary Table 1**). These inbred lines were chosen because each has an available high-quality reference genome ^47^ and accompanying bulk ATAC-seq datasets ^15^, enabling rigorous cross-validation and comparative analysis of regulatory landscapes across genotypes. At this early seedling stage, the hybrids already exhibited pronounced heterotic phenotypes (Student’s t-test, *P* < 0.05; **Supplementary Fig. 1 and Supplementary Table 2**), providing a relevant developmental context for interrogating regulatory mechanisms underlying heterosis. Multiple metrics supported the high quality of the scifi-ATAC-seq data, including transcription start site (TSS) enrichment, ACR summit enrichment, fragment size periodicity, replicate concordance, and consistency with previous scATAC-seq ^32^ and bulk ATAC-seq data ^27,48^ (**Supplementary Fig. 2a–g**). After stringent quality control filtering and genotype demultiplexing, we retained 131,890 single-cell chromatin accessibility profiles, with a median of 2,656 unique Tn5 transposase integrations per nucleus (**Supplementary Fig. 2h, Supplementary Fig. 3 and Supplementary Table 3**).

**Fig. 1.**
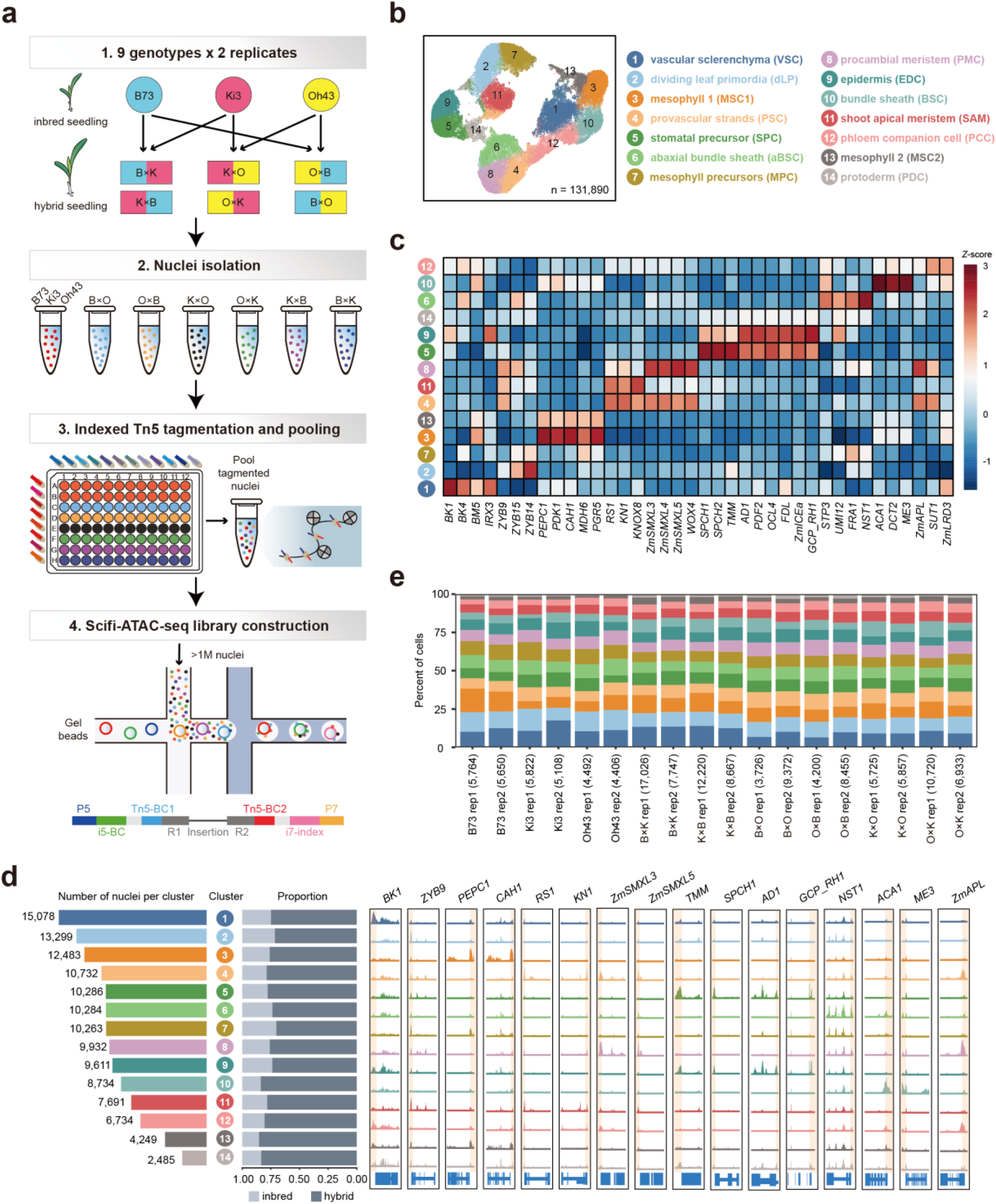
| Single-cell chromatin accessibility landscape of maize. **a**, Schematic of the experimental workflow. **b**, UMAP visualization of maize single-cell chromatin accessibility, colored by major cell clusters. **c**, *Z*-score heatmap of gene chromatin accessibility for marker genes. **d**, Pseudo-bulk chromatin accessibility profiles of the 14 major cell clusters at representative marker genes. Orange shading denotes promoter regions. For each cluster, the total number of nuclei and the relative proportions of inbred and hybrid nuclei are indicated on the left. **e**, Cell-type composition across samples. Colors correspond to cell-type annotations in the UMAPs shown in (**b**).

To define cell identities, we performed unsupervised clustering of nuclei based on genome-wide chromatin accessibility measured at 500-bp resolution. In total, we identified 14 major clusters free of technical variation and with highly consistent nuclei proportions across biological replicates (**Fig. 1b and Supplementary Fig. 4**). Cell-type annotations were performed using multiple complementary approaches, including manual inspection of known marker gene activity, cell-cycle phase inference, and evaluation of enriched biological processes (Methods; **Supplementary Table 4–6**). Using these criteria, each of the 14 clusters was assigned a cell-type identity consistent with a prior study (**Fig. 1b–d and Supplementary Fig. 5–6)** ^33^. We hypothesized that differences in cell-type composition may contribute to organismal heterosis. However, inbreds and hybrids exhibited highly similar cell-type compositions (equivalence test, FDR < 0.05; **Fig. 1e and Supplementary Fig. 4b**), indicating that heterosis is unlikely to be primarily explained by changes in cellular composition.

### Haplotype divergence defines distinct ACR classes and TF motif landscapes

To characterize ACR landscapes across parental inbreds and their hybrids, we first assessed whether hybridization broadly altered global ACR features. For this initial genome-wide comparison, genotype-level pseudo-bulks were generated by pooling reads across cell types and aligned to the B73 v5 reference genome (AGPv5) ^47^, filtered with *WASP* (v0.3.4) ^49^ to reduce reference-mapping bias, downsampled to matched sequencing depths and used for ACR identification. Hybrids showed no consistent changes in ACR number or length relative to inbred lines (permutation test, *P* > 0.05; **Supplementary Fig. 7a–c**). Re-analysis of published bulk ATAC-seq data ^15^ similarly showed no broad hybrid-associated expansion or contraction of accessible chromatin (permutation test, *P* > 0.05; **Supplementary Fig. 7d–f**). To enable accurate haplotype-specific comparisons, we then established haplotype-resolved ACR maps, which we use for all subsequent analyses. Specifically, reads from each genotype × cell type × replicate were aligned to the corresponding parental reference genomes for inbred lines, or to a synthetic diploid assembly generated by concatenating the two parental assemblies for hybrids. ACRs were called independently for each combination, and reproducible ACRs were identified across biological replicates within each cell type before being merged across cell types to generate a non-overlapping ACR set for each inbred genotype or hybrid haplotype. This yielded 43,312– 61,856 ACRs per inbred or hybrid haplotype, with comparable numbers between the two haplotypes of each hybrid (**Fig. 2a**). ACR-to-nearest-gene distance distributions were also highly similar across genotypes, indicating broadly comparable genomic positioning of accessible regions (**Fig. 2b**). We therefore focused on divergence at individual ACRs between the two parental haplotypes. For each parental cross set (for example, the B73–Ki3 set comprising B73, Ki3, B×K and K×B), haplotype-resolved ACRs were classified according to sequence conservation and haplotype-specific chromatin accessible status (Methods; **Fig. 2c**). Sequence-based comparisons revealed that more than 75% of ACRs were conserved between parents, including directly-mapped and minor-shifted ACRs (**Fig. 2d**). In contrast, approximately 5–6% of ACRs were classified as insertion-shifted (IS), comprising small-and large-insertion ACRs (**Supplementary Fig. 8**), 3–4% represented presence-absence variation (PAV) and the remaining ∼15% of ACRs were classified as ambiguous ACRs, as they could not be confidently assigned to a specific category, and were not considered in subsequent analyses (**Fig. 2d**). We further examined chromatin accessible status across haplotypes, classifying ACRs as either shared or variable. The proportion of ACRs with shared accessible status varied markedly across sequence conservation categories, decreasing from over 75% for directly-mapped ACRs to ∼20% for large-insertion ACRs, indicating that structural disruption is associated with increased variability in chromatin accessibility (binomial logistic regression, *P* < 2.2 × 10^-16^; **Fig. 2e**). Together, integration of sequence conservation and chromatin accessibility status allowed ACRs to be grouped into five major categories: conserved-shared, conserved-variable, IS-shared, IS-variable and PAV. Investigating the genomic distributions of ACR categories revealed that conserved-shared and conserved-variable ACRs were strongly enriched at TSSs and gene bodies, respectively (**Fig. 2f**). In contrast, IS and PAV ACRs (collectively, structurally divergent ACR) were enriched in intergenic regions and depleted at TSSs (Fisher’s exact test, FDR < 0.05; **Fig. 2f and Supplementary Fig. 9a–b**). This pattern aligns with prior data indicating that many genome-wide association studies (GWAS) signals fall outside genic regions ^50^, providing a link between structurally divergent ACRs and phenotypic diversity. Indeed, structurally divergent non-genic ACRs were more enriched for GWAS-associated variants ^50^ than genic or conserved non-genic ACRs (Fisher’s exact test, *P* < 0.05; **Fig. 2g**), with a substantial subset (>73.9%) also harboring at least one expression quantitative trait locus (eQTL) signal ^51^.

**Fig. 2.**
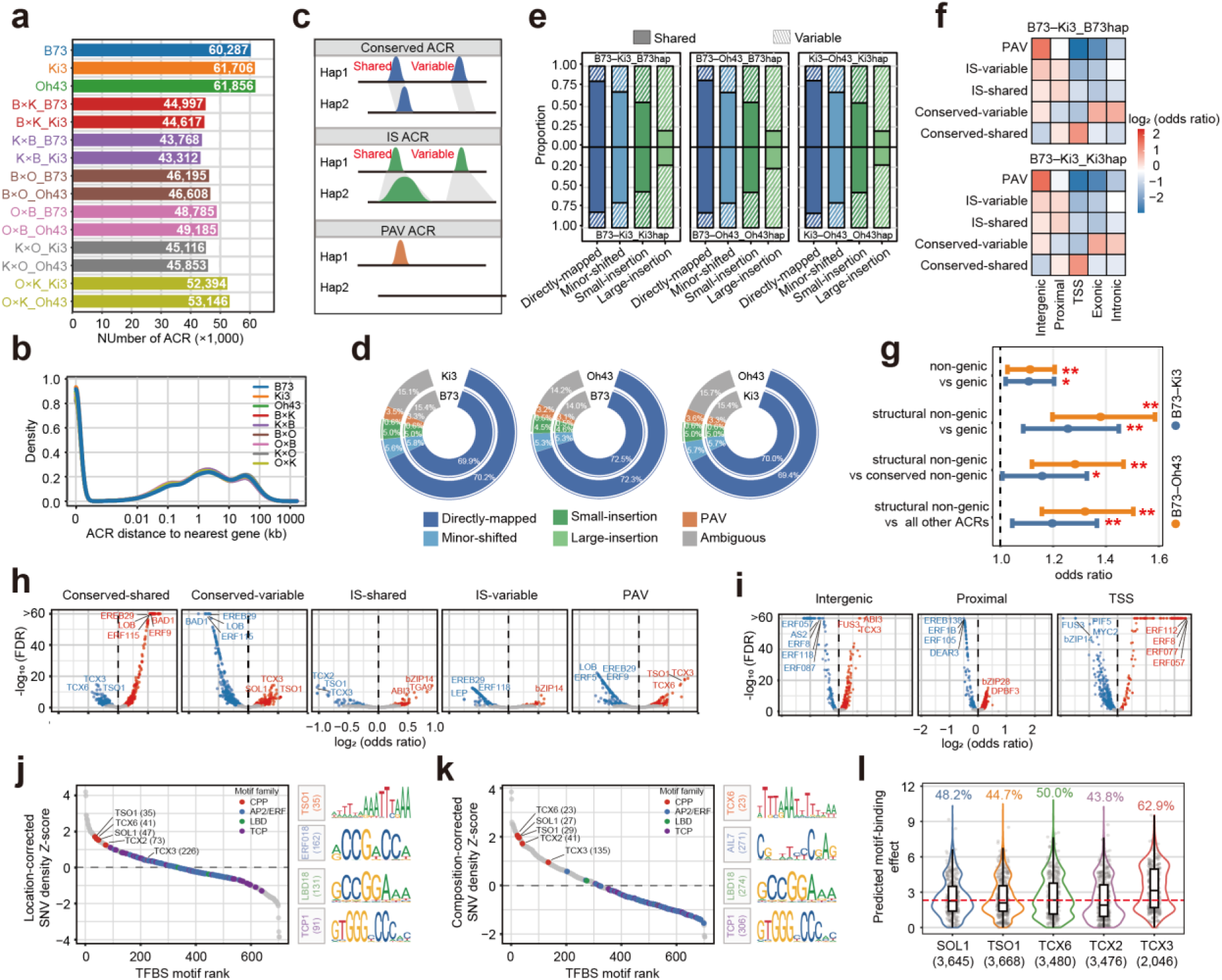
| Characterization of ACRs. **a**, Number of ACRs identified in each genotype or haplotype. **b**, Density distributions of ACR–gene distances across genotypes. **c**, Schematic overview of ACR classification based on sequence conservation and accessibility status. Colored peaks indicate chromatin accessibility signal. Gray shading indicates conserved or alignable sequence between haplotypes; no shading indicates sequence absence or no clear alignable counterpart. **d**, Proportional composition of ACRs across sequence-based categories in each haplotype. **e**, Proportions of shared and variable ACRs across sequence-based categories for each parental–hybrid combination. **f**, Enrichment of ACR categories across genomic annotation classes for each haplotype. **g**, Enrichment of GWAS-associated variants across ACR categories. Odds ratios (points) and 95% confidence intervals (lines) for B73–Ki3 (blue) and B73–Oh43 (orange). Two-sided Fisher’s exact test was used to determine statistical significance (\**P* < 0.05, \*\**P* < 0.01). **h**, TF binding motif enrichment and depletion across sequence/accessibility-defined ACR categories in the B73–Ki3 parental–hybrid combination (B73, Ki3, B×K and K×B). **i**, TF binding motif enrichment and depletion across ACRs stratified by genomic context for the B73–Ki3 parental–hybrid combination. **j**, TF binding motifs ranked by local-background-corrected SNV density Z-score within sequence-conserved ACRs. Colors denote CPP, AP2/ERF, LBD and TCP motifs; sequence logos show the highest-ranked motif from each family. Numbers in parentheses indicate motif rank. **k**, As in (**j**) after correction for trinucleotide sequence context. **l**, Distributions show the absolute *PERFECTOS-APE*-predicted binding effects for polymorphic CPP motif–SNV pairs. The dashed red line marks a fivefold allele difference; percentages and numbers indicate the fraction and number of scored motif–SNV pairs, respectively.

We next asked whether these distinct ACR categories exhibit differential TF binding motif preferences. Conserved-shared ACRs were strongly enriched for motifs corresponding to core developmental TF families, including APETALA2/ETHYLENE RESPONSIVE FACTOR (AP2/ERF), TEOSINTE BRANCHED1/CYCLOIDEA/PROLIFERATING CELL FACTOR (TCP) and LATERAL ORGAN BOUNDARY DOMAIN (LBD). In contrast, variable ACR categories showed pronounced enrichment for CYSTEINE-RICH POLYCOMB-LIKE PROTEIN (CPP) family, previously found associated with developmental plasticity, cell proliferation and stress-responsive regulation ^52,53^ (Fisher’s exact test, FDR < 0.05; **Fig. 2h and Supplementary Fig. 9c**). Notably, TF binding motifs enriched in conserved-shared ACRs were depleted in variable ACR categories, and *vice versa*, revealing a reciprocal pattern of TF binding motif enrichment driven by chromatin accessibility variance. Restricting the analysis to non-genic ACRs to minimize gene-body sequence constraints, stratification by genomic context revealed a similar trend, consistent with previous reports ^15^ and mirrored the motif enrichment patterns observed above; TSS ACRs were preferentially enriched for AP2/ERF motifs, intergenic ACRs were enrichment for CPP motifs, and proximal ACRs exhibited intermediate profiles (Fisher’s exact test, FDR < 0.05; **Fig. 2i and Supplementary Fig. 9d**). Thus, TF binding motif was partitioned along a regulatory-context gradient, with conserved-shared and TSS-associated ACRs enriched for motifs linked to core developmental regulation, whereas variable and intergenic ACRs were enriched for motif families associated with cell proliferation, potentially reflecting differences in developmental velocity.

Given the distinct TF binding motif enrichments across ACR categories with differing sequence constraints, we examined whether the TF binding motifs also differed in sequence polymorphism. To minimize confounding effects from large-scale structural variation, we restricted the analysis to sequence-conserved ACRs (∼75% of total ACRs; **Fig. 2d**) and quantified single-nucleotide variant (SNV) density within motifs. CPP motifs remained highly polymorphic after correcting for the local SNV density of their host ACRs, ranking above most AP2/ERF, TCP and LBD motifs (**Fig. 2j and Supplementary Table 7**). This pattern persisted after accounting for trinucleotide sequence context (**Fig. 2k and Supplementary Table 8**), indicating that the elevated sequence variation at CPP motif sites could not be explained solely by their local regulatory environment or motif sequence composition. To assess the potential consequences of this variation, we predicted allele-specific effects on TF binding using *PERFECTOS-APE* ^54^, with 43.8–62.9% of polymorphic motif–SNV pairs across the five CPP motifs predicted to substantially alter TF-binding affinity (Methods; **Fig. 2l**). Together, these results identify CPP motif sites as a particularly polymorphic class of regulatory sequence, potentially providing a substrate for regulatory variation in CPP-associated cell proliferation and early seedling development.

### Hybridization attenuates chromatin accessibility cell-type specificity

Having defined ACR categories with distinct genomic distributions and TF motif profiles, we next asked whether these categories also differed in cell-type specificity. To this end, we first identified cell-type-specific ACRs (ctACRs), characterized by accessibility concentrated in a limited number of cell types (Methods; **Fig. 3a**). Compared to non-cell-type-specific ACRs (nctACRs), ctACRs were more likely to fall within accessible-status-variable categories, with the highest proportion observed in conserved-variable ACRs (Fisher’s exact test, FDR < 0.05; **Fig. 3b and Supplementary Fig. 10a–b**). Cell-type specificity also varied across genomic contexts, with TSS and proximal ACRs being predominantly non-cell-type-specific, whereas ctACRs were enriched in distal regions (Fisher’s exact test, FDR < 0.05; **Fig. 3c and Supplementary Fig. 10c–d**), consistent with other plant species with large genomes ^39^. Interestingly, cell-type specificity was also prominent among genic ACRs (Fisher’s exact test, FDR < 0.05; **Fig. 3c and Supplementary Fig. 10c–d**). As gene-body accessibility may reflect a transcription-permissive chromatin architecture ^55^, we asked whether the cell-type specificity of genic ACRs could be partly explained by cell-type-restricted transcription. Analysis of previously published maize single-nucleus RNA-seq data ^33^ showed that genes linked to genic ctACRs were themselves more cell-type-specific in transcription than genes linked to genic nctACRs (Wilcoxon rank sum test, *P* < 0.05; **Supplementary Fig. 10e–f**). Thus, the elevated cell-type specificity among genic ACRs likely reflects, at least in part, transcription-associated chromatin accessibility.

**Fig. 3.**
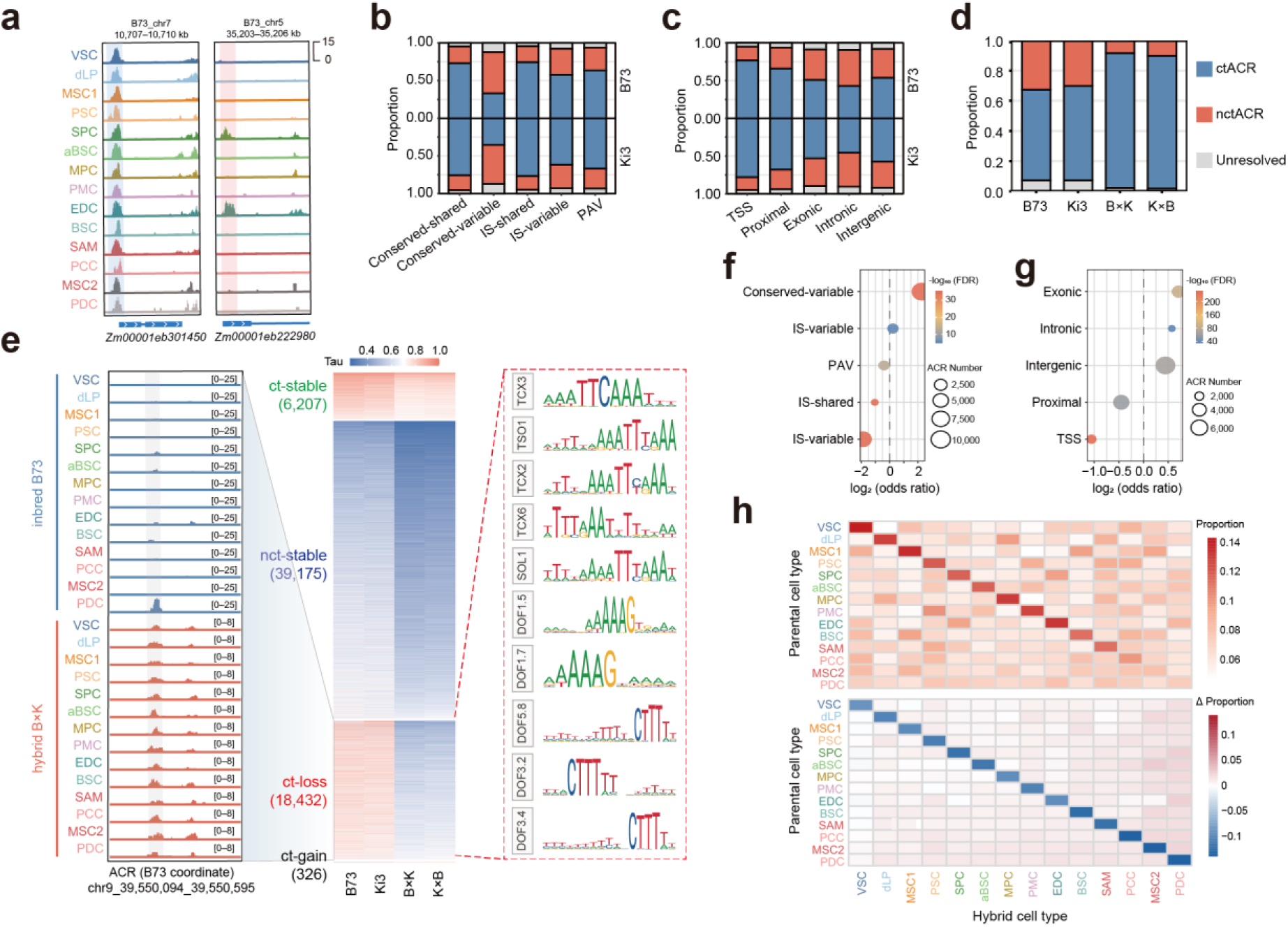
| Hybridization attenuates chromatin accessibility cell-type specificity. **a**, Genome browser tracks illustrating the examples of ctACR (left) and nctACR (right). Shaded regions mark the ACR region**. b**, ctACR and nctACR proportions across sequence/accessibility-defined ACR categories, shown separately for the two parents. **c**, ctACR and nctACR proportions across genomic contexts, shown separately for the two parents. **d**, ctACR and nctACR proportions in B73, Ki3 and their reciprocal hybrids B×K and K×B. **e**, Tau heatmap across B73, Ki3 and reciprocal hybrids showing four ACR specificity-state groups. Left, representative ct-loss ACR showing attenuation of chromatin accessibility cell-type specificity in B×K relative to B73. Right, representative TF binding motifs enriched in ct-loss ACRs (FDR < 0.05, odds ratio > 1.5). Shaded regions mark the ACR region. **f**, Enrichment of sequence/accessibility-defined ACR categories for ct-loss ACRs. **g**, Enrichment of genomic contexts for ct-loss ACRs. **h**, Hybrid accessibility profiles of ct-loss ACRs grouped by parental source cell type, defined as the cell type with the highest mean accessibility proportion across two parents. Top heatmap, mean accessibility proportion across hybrid cell types. Bottom heatmap, Δ proportion between hybrids and parents. Accessibility proportion was calculated for each ACR as the fraction of CPM-normalized accessibility assigned to each cell type across all 14 cell types and Δ proportion denotes hybrid mean minus parental mean proportion.

With ctACRs defined in parental inbreds, we next examined whether hybridization alters the overall cell-type specificity of chromatin accessibility. Approximately 30–40% of parental ACRs were cell-type-specific, whereas this fraction was significantly reduced in hybrids, with a reciprocal increase in nctACRs (**Fig. 3d and Supplementary Fig. 10g–h**). Considering Tau-based of cell-type specificity can be sensitive to sequencing depth, we tested whether the reduced specificity in hybrids could be explained by their greater sequencing depth (Methods). We therefore downsampled inbred and hybrid datasets to matched sequencing depths and re-calculated chromatin accessibility cell-type specificity. Although Tau values varied with sequencing depth, hybrids consistently retained lower cell-type specificity than inbreds at equivalent coverage (Wilcoxon rank sum test, *P* < 2.2 × 10^-16^; **Supplementary Fig. 10i**), indicating that hybridization attenuates cell-type-specific chromatin accessibility beyond differences in sequencing depth. To ask whether attenuated chromatin accessibility cell-type specificity in hybrids is also reflected at the transcriptional level, we analyzed an independent maize developmental transcriptome dataset spanning 23 tissues in parental inbreds and hybrids ^56^. Although these data represent tissue-level rather than cell-type-level variation, they provide an orthogonal framework to test whether hybridization is associated with attenuated context-specific gene activity. Consistent with the chromatin accessibility results, hybrids showed a similar attenuation in expression specificity across tissues compared with parental inbreds (Wilcoxon rank sum test, *P* < 1.8 × 10^-5^; **Supplementary Fig. 10j**). Together, these analyses indicate that hybridization is associated with a broader deployment of regulatory activity across cellular contexts.

To identify the regulatory elements affected by attenuated cell-type specificity, we classified ACRs according to whether their cell-type specificity was maintained or altered between parental inbreds and hybrids. This resolved four specificity-state groups, representing ACRs that remained cell-type-specific in both states (ct-stable), were broadly accessible in both states (nct-stable), shifted from parental cell-type-specific to broadly accessible in hybrids, with a median decrease in Tau of at least 0.15 across all three crosses (ct-loss), or gained cell-type specificity in hybrids (ct-gain) (**Fig. 3e and Supplementary Fig. 10k–m**). The nct-stable class predominated, whereas ct-gain ACRs were rare and were not considered further. Because ct-loss ACRs directly capture the hybrid-associated attenuation of parental cell-type specificity, we focused subsequent analyses on this class. CPP and DOF family motifs were significantly enriched in ct-loss ACRs (Fisher’s exact test, FDR < 0.05, odds ratio > 1.5; **Fig. 3e and Supplementary Fig. 10k–l**). ct-loss ACRs were also enriched in the conserved-variable and intergenic categories (Fisher’s exact test, FDR < 0.05, **Fig. 3f–g and Supplementary Fig. 10n–o**), consistent with our earlier observations (Fisher’s exact test, FDR < 0.05, **Fig. 2h–i and Supplementary Fig. 9c–d**). Moreover, genes nearest to ct-loss ACRs showed the largest decrease in expression tissue-specificity among the four groups (Permutation test, *P* = 0.009; **Supplementary Fig. 10p**). Together, these results link attenuated specificity in hybrids to a CPP-and DOF-marked subset of distal, conserved-variable ACRs and suggest that chromatin-level attenuation propagates to nearby gene expression.

We next quantified the cellular breadth of individual ct-loss ACRs by counting the number of cell types classified as accessible using a two-state negative-binomial mixture model. ct-loss ACRs were accessible across more cell types in hybrids than in the parental inbreds, with a median increase in accessibility breadth of 3 cell types, revealing an expanded breadth of chromatin accessibility at individual loci (**Supplementary Fig. 10q**). We then asked whether this broadening reflected consistent redirection towards particular alternative cell identities. After grouping ct-loss ACRs by parental source cell type, defined as the cell type with the highest mean accessibility proportion across the two parents, we found that hybrid chromatin accessibility remained highest in the corresponding cell-type dimension. However, we found that source-cell bias was reduced, and no alternative cell type consistently became dominant (**Fig. 3h and Supplementary Fig. 10r–s**). Thus, chromatin accessibility attenuation effect in hybrids weakens parental cell-type bias while distributing relative accessibility more broadly across all cell types.

### Hybrid chromatin inheritance is shaped by parental divergence and celluar context

To establish the parental regulatory differences inherited by hybrids, we quantified chromatin accessibility divergence between parental inbreds across cell types. We focused on putative regulatory ACRs by excluding exonic intervals and identified differentially accessible regions (DARs) between parental lines within each cell type (fold change > 2; FDR < 0.05). An average of 2,065 DARs per cell type (range: 498–3,512) were detected, spanning divergence magnitudes from moderate (2–4-fold) to strong (>4-fold) accessibility differences, as well as single-parent ACRs (SPA), which represent the most extreme form of parental bias (**Fig. 4a and Supplementary Fig. 11a–b**). DAR counts were directionally skewed toward one parent in most cell types, indicating asymmetric chromatin accessibility between parental genomes (**Fig. 4a and Supplementary Fig. 11a–b**). Despite this range of divergence magnitudes, both DARs and SPAs exhibited similar distribution patterns across cell types (**Fig. 4b–c and Supplementary Fig. 11c–d**). The majority (46–69%) were confined to fewer than 25% of cell types, whereas only a small fraction (6–14%) were broadly shared across more than 75% of cell types. (**Fig. 4b and Supplementary Fig. 11c–d**). Stratification by directional consistency further illustrated that most parent-biased DARs and SPAs maintained the same bias direction across cell types, whereas mixed-bias cases were nearly absent. (**Fig. 4d and Supplementary Fig. 11e–f**). Thus, parental chromatin accessibility divergence is largely cell-type-restricted and directionally stable across cellular contexts.

**Fig. 4.**
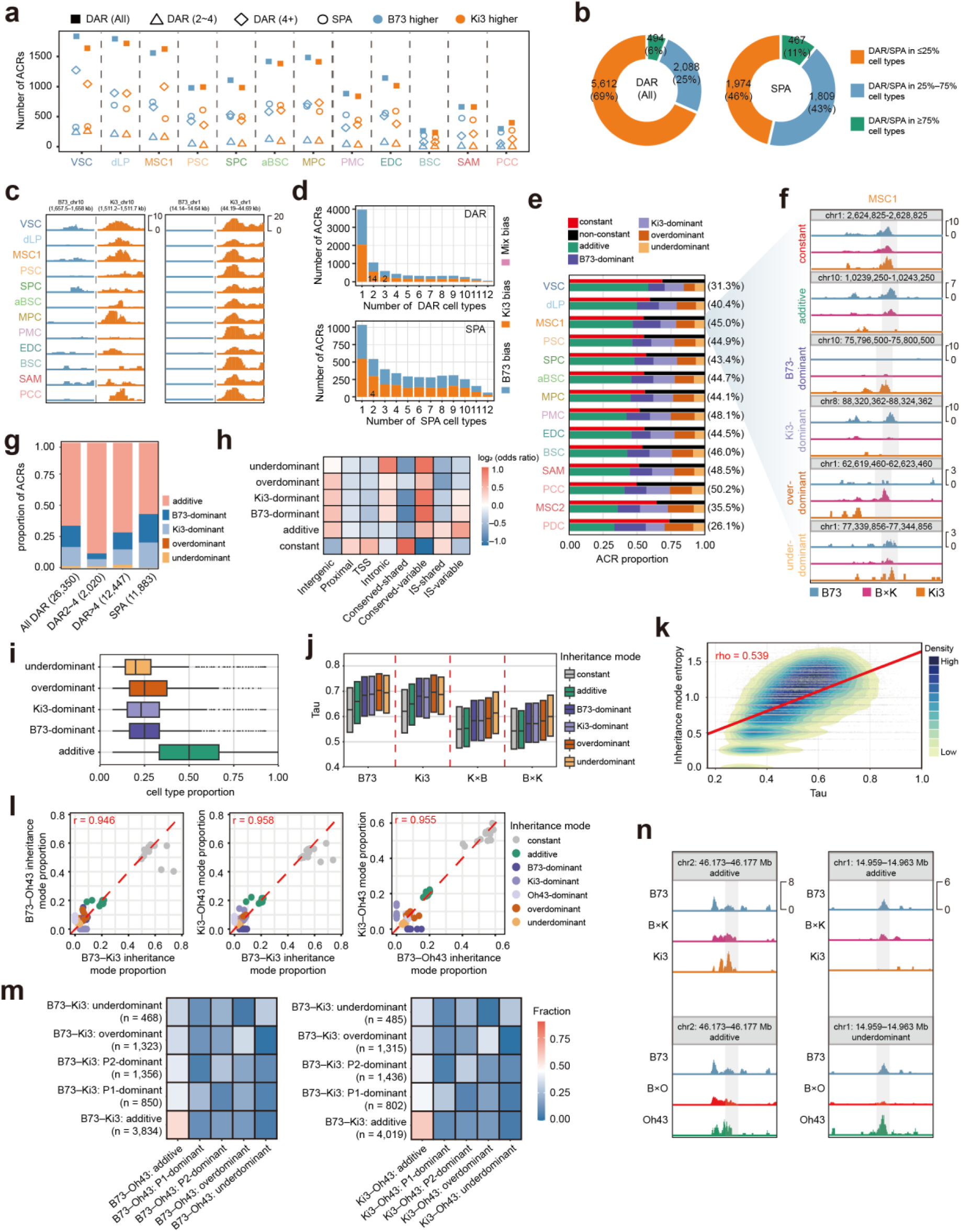
| Cell-type-resolved parental divergence and chromatin inheritance in maize hybrids. **a**, The number of DARs in each cell type is shown, with parental bias indicated by color. MSC2 and PDC cell types are excluded because they have too few DARs for meaningful visualization. **b**, DARs and SPAs were grouped by cell-type recurrence, defined as regions detected in ≤25%, 25%–75%, or ≥75% of tested cell types. Donut plots show the distribution of DARs (left) and SPAs (right) across these categories. **c**, Representative genome browser tracks illustrating two modes of parental accessibility divergence: divergence restricted to a subset of cell types (left) or consistently observed across all cell types (right). **d**, Numbers of DARs (top) and SPAs (bottom) detected across increasing numbers of cell types. Colors indicate the directional consistency of parental accessibility divergence, with regions showing consistent B73 bias, consistent Ki3 bias, or mixed bias across cell types. Counts of ACRs with mixed bias are annotated on bars, unlabeled bars indicate zero. **e**, Cell-type-specific composition of ACR inheritance modes in the B×K hybrid comparison. For each cell type, the upper bar shows the proportion of constant versus non-constant ACRs, and the lower bar shows the relative composition of non-constant ACRs across additive, parental-dominant, overdominant, and underdominant categories. Numbers in parentheses indicate the proportion of non-constant ACRs in each cell type. **f**, Representative genome browser views illustrating distinct ACR patterns in MSC1 cell type. Tracks show accessibility signals from the two parental inbreds (B73 and Ki3) and one of the hybrids (B×K), highlighting examples of constant, additive, B73-dominant, Ki3-dominant, overdominant, and underdominant ACRs. Shaded regions mark the representative ACRs. **g**, Stacked bar plots show the proportions of ACRs exhibiting distinct inheritance patterns within each parental DAR class. Numbers in parentheses indicate the total number of DARs summed across cell types. **h**, Heat map showing enrichment of ACR inheritance modes across genomic contexts and sequence conservation/accessibility classes. **i**, The set of ACRs assigned to each non-constant inheritance mode was used to examine the breadth of inheritance patterns across cell types. For each ACR–mode pair, we determined the fraction of cell types in which the inheritance mode was observed and used boxplots to visualize the distribution of values. **j**, Distribution of Tau for ACRs grouped by inheritance mode in B73, Ki3 and reciprocal hybrids, pooled across cell types. Pairwise differences between inheritance modes within each genotype were assessed using two-sided Wilcoxon rank-sum tests. **k**, Density plot showing the association between Tau (mean of reciprocal hybrids) and Shannon entropy of inheritance modes across all cell types. Colors indicate local point density. Spearman’s rho is indicated. **l**, Global similarity of inheritance-mode composition across hybrids. Pairwise comparisons between B73–Ki3, B73–Oh43 and Ki3–Oh43 are shown. Each point represents the proportion of ACRs assigned to a given inheritance mode within a cell type. Pearson correlation coefficients were calculated and shown. **m**, Aggregated switching rate matrices of chromatin accessibility inheritance modes across hybrid combinations after pooling all cell types. Rows indicate modes in the reference hybrid and columns indicate modes in the comparison hybrid. (left) B73–Ki3 vs B73–Oh43. (right) B73–Ki3 vs Ki3–Oh43. Colors denote switching rate. Numbers in parentheses indicate the number of loci assigned to each row category. In dominant categories, P1 denotes the parent shared between the two hybrids (e.g., B73 in B73–Ki3 vs B73–Oh43), whereas P2 denotes the parent that differs between the hybrids (e.g., Ki3 in B73–Ki3 and Oh43 in B73–Oh43). **n**, Representative loci illustrating conserved and switching inheritance modes across hybrid combinations. Chromatin accessibility tracks for parental lines and corresponding hybrids are shown. (left) Example of a locus with conserved additive inheritance across hybrids. (right) Example of a locus exhibiting a transition in inheritance mode between hybrids.

We next examined how chromatin accessibility is inherited in hybrid genomes. As we did not observe parent-of-origin effects (**Supplementary Fig. 12**), we treated reciprocal hybrids jointly in subsequent inheritance analyses. ACRs were then classified into distinct inheritance modes for each cell type, including constant, additive, parental-dominant, overdominant and underdominant categories. Across hybrid combinations and cell types, 40.3–73.9% of ACRs were constant. Among variable ACRs, additive inheritance predominated (∼40–60%), whereas non-additive modes were collectively more frequent than in bulk analyses ^15,16^, with parental dominance, overdominance and underdominance averaging 29.4%, 14.4% and 7.1%, respectively (**Fig. 4e–f and Supplementary Fig. 13a–b**). Stratification by parental DAR levels showed that additive inheritance contributed across all divergence classes, but its relative contribution was highest among moderate-effect DARs (2–4-fold). Strongly divergent DARs and SPAs (>4-fold) showed a corresponding increase in parental-dominant inheritance, indicating a modest shift towards non-additive chromatin accessibility states with increasing parental divergence (**Fig. 4g and Supplementary Fig. 13c–d**). Beyond differences in parental divergence, inheritance modes also exhibited non-random distributions across genomic contexts and sequence/accessibility-defined ACR categories. For example, constant ACRs were strongly enriched in proximal and TSS-associated regions and preferentially associated with shared-accessible-status categories, whereas non-constant ACRs showed the opposite pattern, as expected (Fisher’s exact test, FDR < 0.05; **Fig. 4h and Supplementary Fig. 13e–f**) ^15,16^.

We further assessed the cell-type breadth of inheritance modes at individual ACRs. Additive inheritance showed broader chromatin accessibility across cell types, whereas non-additive ACRs exhibited greater cell-type specificity (Wilcoxon rank sum test, *P* < 2.2 × 10^-16^; **Fig. 4I–J and Supplementary Fig. 13g–j**). Moreover, inheritance-mode entropy increased with cell-type specificity, indicating that ctACRs tend to exhibit more variable inheritance outcomes (Spearman’s rho > 0.5; **Fig. 4k and Supplementary Fig. 13k–l**). Together, these results indicate that non-additive chromatin inheritance is closely associated with cell-type-dependent regulatory programs. Next, we asked whether inheritance modes are conserved across the three different hybrid combinations. To address this, we compared inheritance-mode composition between hybrids sharing one parental genotype. Despite differences in the second parental background, inheritance-mode compositions were highly similar across hybrid combinations, with strong correlations in inheritance mode proportions across cell types (Pearson’s r > 0.94; **Fig. 4l**). At individual ACR level, inheritance modes were partially conserved across different hybrids. Additive loci showed the strongest inheritance mode retention. However, ACRs in all inheritance categories exhibited hybrid-dependent switching (**Fig. 4m–n and Supplementary Fig. 14**). ACRs with higher switching rates were enriched near genes involved in translation and ribosome-related processes (GSEA analysis, *P* < 0.05; **Supplementary Table 9**), suggesting that hybrid-dependent inheritance switching is biased toward regulatory neighborhoods of protein biosynthesis processes. Thus, global inheritance-mode composition is broadly conserved across hybrids, whereas individual loci can undergo functionally biased, hybrid-dependent switching.

### *Cis-* and *trans-*acting effects underlie hybrid chromatin divergence

Allele-biased chromatin accessibility has been observed in hybrids across diverse species, including maize ^13–17^. By leveraging our scifi-ATAC-seq data, we were able to investigate the cell-type-resolved regulatory basis of chromatin accessibility divergence in maize hybrids and partitioned this divergence into *cis*-and *trans*-regulatory components. Parental differences capture the combined effects of *cis*-and *trans*-acting variation, whereas allelic imbalance within hybrids provides a readout of *cis* effects under a shared *trans* environment ^57^. We applied a maximum likelihood framework to assign ACRs to non-divergent, *cis*-only, *trans*-only and *cis* & *trans* categories, with the latter further separated by the direction and relative strength of the two regulatory components (**Fig. 5a and Supplementary Fig. 15a**). For example, ACRs near *MTERF-DOMAIN PROTEIN1* (*MTERF1*), *SBP-TRANSCRIPTION FACTOR 3* (*SBP3*) and *INDETERMINATE GAMETOPHYTE1* (*IG1*) illustrated these allele-resolved patterns within representative cell types, corresponding to *cis*-only, *trans*-only and *cis* & *trans* effects, respectively (**Fig. 5b**). Quantitatively, non-divergent ACRs accounted for 50.0–61.0% of ACRs across hybrids and cell types, indicating that most ACRs showed no detectable parental chromatin accessibility divergence or allelic imbalance in hybrids, broadly aligning with previous findings of gene expression in mice ^58,59^. Among the divergent categories, *cis*-only effects represented the largest class, accounting for 24.8% of ACRs on average, whereas *trans*-only and *cis* & *trans* categories accounted for 12.2% and 7.7%, respectively. Among *cis* & *trans* ACRs, compensating configurations (opposite-direction *cis* and *trans* effects) were more common than reinforcing configurations (same-direction effects), accounting for ∼60% of cases. Across both configurations, *trans*-dominant subclasses, in which the inferred *trans* component exceeded the *cis* component, comprised more than 75% of *cis* & *trans* ACRs (**Fig. 5a and Supplementary Fig. 15a**). Together, *cis*-only effects formed the largest divergent class overall, whereas *trans*-regulatory effects were a consistent and prominent feature of loci shaped jointly by *cis*-and *trans*-regulatory variation.

**Fig. 5.**
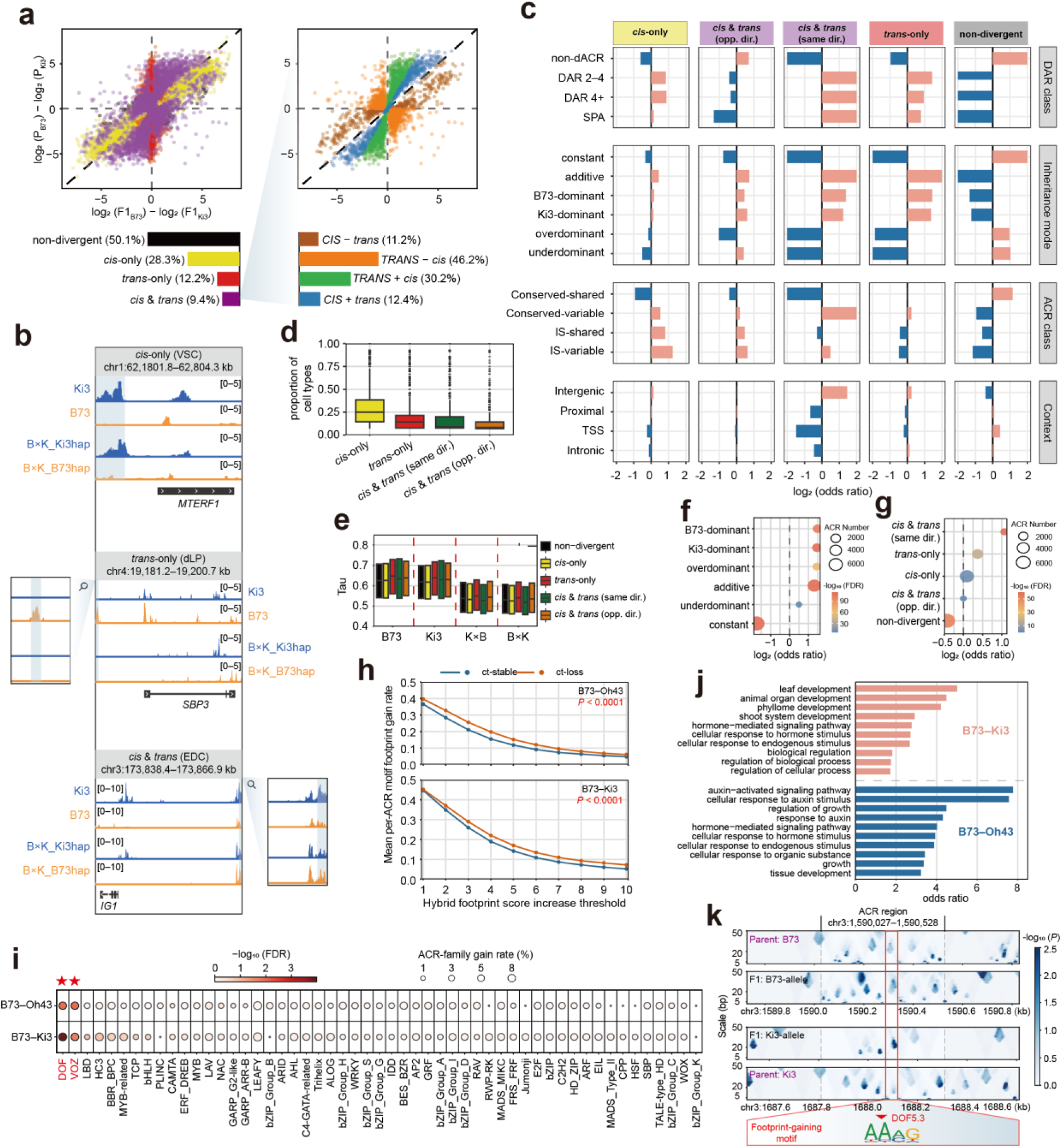
| Cell-type-resolved regulatory divergence and TF footprint remodeling in maize hybrids. **a**, *Cis*-and *trans*-regulatory inference for ACRs in the VSC cell type of B73–Ki3. Left, classification of ACRs into non-divergent, *cis*-only, *trans*-only and *cis* & *trans* categories. Right, subclassification of *cis* & *trans* ACRs by the direction and relative strength of *cis*-and *trans*-regulatory effects. Percentages indicate category proportions. **b**, Representative haplotype-resolved accessibility profiles illustrating *cis*-only, *trans*-only and *cis* & *trans* effects at the *MTERF1*, *SBP3* and *IG1* loci, respectively. Shaded regions mark the representative ACRs and magnified views of the *SBP3* and *IG1* ACRs are shown alongside the corresponding loci. **c**, Enrichment of regulatory categories across parental DAR classes, inheritance modes, ACR sequence/accessibility classes and genomic contexts in B73–Ki3. Bars show log_2_ odds ratios from Fisher’s exact tests; values beyond ±2 were capped at ±2 for visualization. Red and blue indicate significant enrichment and depletion, respectively (FDR < 0.05), and grey indicates non-significant associations. **d**, Cell-type recurrence of regulatory effects in B73–Ki3. For each ACR, recurrence was calculated as the fraction of cell types in which it was assigned to each category. **e**, Tau scores of B73–Ki3 ACRs assigned to each category across parental and hybrid genotypes. **f**, Enrichment of ct-loss ACRs across inheritance mode categories in B73– Ki3. **g**, Enrichment of ct-loss ACRs across *cis*/*trans* effect categories in B73–Ki3. **h**, Mean per-ACR fraction of motifs showing parent-to-hybrid footprint gains across increasing footprint-score thresholds in ct-stable and ct-loss ACRs. *P* values are from one-sided permutation tests comparing ct-loss with ct-stable. **i**, Motif-family enrichment among high-confidence footprint-gaining events in ct-loss ACRs. Red asterisks mark families significantly enriched in both crosses. **j**, GO enrichment for genes linked to footprint-gaining ct-loss ACRs. Bars show odds ratios for the top enriched biological processes in B73–Ki3 and B73–Oh43. **k**, Representative view of a recurrent DOF-family footprint gain at a ct-loss ACR within the *GLK2* locus in EDC cells.

We next asked how these *cis*-and *trans*-regulatory categories relate to hybrid inheritance modes and broader chromatin features. As expected, non-divergent ACRs were significantly enriched among non-dACRs, constant ACRs and conserved-shared ACRs. Divergent ACRs were instead enriched among additive and parental-dominant inheritance modes, broadly consistent with previous maize bulk-ATAC analyses of chromatin inheritance (Fisher’s exact test, FDR < 0.05; **Fig. 5c and Supplementary Fig. 15b–c**) ^16^. Stratification by parental divergence revealed clearer differences among regulatory categories. *Cis*-only, *trans*-only and same-direction *cis* & *trans* effects were enriched among parental DARs, linking parental chromatin accessibility divergence to local allele-linked variation, *trans*-acting effects or coordinated *cis*/*trans* effects. Opposite-direction *cis* & *trans* ACRs showed a distinct pattern, with enrichment among non-dACRs and depletion from stronger DAR and SPA classes, consistent with compensating effects buffering parental divergence (Fisher’s exact test, FDR < 0.05; **Fig. 5c and Supplementary Fig. 15b–c**). These *cis*/*trans* regulatory categories also differed across sequence/accessibility classes and genomic contexts. For example, conserved-variable ACRs were broadly associated with regulatory divergence, while IS-associated ACRs preferentially marked *cis*-only and combined *cis* and *trans* effects rather than *trans*-only effects (Fisher’s exact test, FDR < 0.05; **Fig. 5c and Supplementary Fig. 15b–c**). This pattern is consistent with local insertion-associated sequence variation contributing to *cis*-dependent divergence, whereas *trans*-only effects are less dependent on local sequence differences and instead act on sequence-conserved but accessible-status-variable regulatory elements.

Consideration of genomic context indicated that *cis*-only, *trans*-only and same-direction *cis* & *trans* effects were enriched in intergenic regions and depleted near TSSs (Fisher’s exact test, FDR < 0.05; **Fig. 5c and Supplementary Fig. 15b–c**). Given that transposable elements (TEs) are a major source of distal regulatory variation in plant genomes and have been implicated in maize hybrid vigor ^18,19^, this distal bias prompted us to ask whether these categories were preferentially TE-associated. Among intergenic ACRs, TE overlap mirrored the distal enrichment of these *cis*/*trans* regulatory categories, being strongest for same-direction *cis* & *trans* ACRs, weaker but still significant for *cis*-only and *trans*-only ACRs and depleted for non-divergent and opposite-direction *cis* & *trans* ACRs (Fisher’s exact test, *P* < 0.05; **Supplementary Fig. 15d**). This pattern was further resolved at the TE family level, with terminal inverted repeat (TIR) transposon-and Helitron-associated ACRs enriched in *cis*-only and *trans*-only categories, whereas ACRs associated with long terminal repeat (LTR) retrotransposons were enriched in same-direction *cis* & *trans* ACRs (**Supplementary Fig. 15e**). Using TE sequence divergence as a proxy for relative age, we found that LTR-associated same-direction *cis* & *trans* ACRs were biased toward older, more diverged LTR fragments, whereas opposite-direction *cis* & *trans* ACRs were associated with younger LTRs; age differences among TIR-and Helitron-associated ACRs were weak or absent (Wilcoxon rank sum test; **Supplementary Fig. 15f**). Together, these results suggest that TE-associated regulatory divergence is structured by TE family and age, with older LTR fragments preferentially linked to coordinated *cis* & *trans* effects. More broadly, TE families of different evolutionary ages appear to contribute through distinct *cis*/*trans* regulatory architectures to hybrid chromatin accessibility divergence.

### Regulatory features associated with attenuated cell-type specificity

Exploring the role of cell identity towards *cis*/*trans* effects, we asked whether effect classifications were consistent at the same ACRs in different cell-type contexts. *Cis*-only effects were most broadly maintained across cell types, whereas *trans*-only and *cis* & *trans* effects were more restricted (**Fig. 5d and Supplementary Fig. 16a**). Consistent with this restricted deployment, *trans*-only ACRs showed the highest cell-type specificity among categories (Wilcoxon rank sum test, *P* < 2.2 × 10^-16^; **Fig. 5e and Supplementary Fig. 16b**), suggesting that *trans*-regulatory divergence preferentially affects highly cell-type-specific regulatory elements. Given the strong cell-type dependence of regulatory divergence, we focused on ct-loss ACRs to examine the regulatory features associated with hybrid attenuation of cell-type specificity. ct-loss ACRs were depleted from constant loci and enriched across non-constant inheritance modes, particularly non-additive categories (Fisher’s exact test, *P* < 0.05; **Fig. 5f and Supplementary Fig. 16c**), indicating that attenuation is accompanied by altered inheritance modes in hybrids and may be linked to regulatory patterns associated with heterosis. Partitioning ct-loss ACR by *cis*/*trans* category further showed the strongest enrichment for same-direction *cis* & *trans* effects, followed by *trans*-only effects, whereas *cis*-only effects showed little enrichment (Fisher’s exact test, *P* < 0.05; **Fig. 5g and Supplementary Fig. 16d**). These results link hybrid attenuation of cell-type specificity to altered accessibility inheritance and regulatory divergence involving *trans* effects, rather than local *cis* effects alone.

Given the enrichment of *trans*-involved effects in ct-loss ACRs and the close association between chromatin accessibility and TF binding, we asked whether the broadening of accessibility across hybrid cell types was accompanied by gains in TF occupancy. We restricted this analysis to the B73–Ki3 and B73–Oh43 ACR sets, both of which could be anchored directly to a common B73 coordinate system, thereby avoiding the additional cross-assembly projection required for Ki3– Oh43. We quantified parent-to-hybrid footprint gains across ACR–motif pairs, requiring support in at least two cell types, and summarized their prevalence as the fraction of gain motifs within each ACR. Across a range of footprint-score-gaining thresholds, ct-loss ACRs consistently exhibited higher motif-gaining fractions than ct-stable ACRs, which retained high cell-type specificity in hybrids (Permutation test, *P* < 0.0001; **Fig. 5h**). Motif-family analysis further identified DOF and VASCULAR PLANT ONE-ZINC-FINGER (VOZ) as the TF families most enriched among footprint-gaining events across crosses, consistent with our earlier finding of DOF motif enrichment specific to ct-loss ACRs (Fisher’s exact test, FDR < 0.05; **Fig. 5i**). Genes linked to gain-bearing ct-loss ACRs were enriched for developmental and hormone-related processes, including leaf and shoot development, hormone signaling and growth (Fisher’s exact test, FDR < 0.05; **Fig. 5j and Supplementary Table 10**), suggesting that footprint gains are preferentially concentrated near developmental genes. Notably, an intronic ct-loss ACR within *GOLDEN2-like* (*GLK2*) showed DOF-family footprint gains across multiple hybrid cell types, particularly in dLP, aBSC, MPC and EDC, where most DOF motifs exhibited increased footprint scores across the B73–Ki3 and B73–Oh43 crosses(**Fig. 5k and Supplementary Fig. 16e–f**). *GLK2* encodes a canonical regulator of bundle-sheath chloroplast differentiation and contributes to the cellular specialization of C4 photosynthesis in plants ^60^. In parallel, comparative analyses of rice and sorghum have shown that DOF motifs form part of a conserved bundle-sheath *cis*-regulatory code and are recruited by photosynthetic genes during the evolution of C4 bundle-sheath expression ^41^. Although our data does not identify the responsible DOF factor or establish direct DOF regulation of *GLK2*, this locus provides an example of hybrid-associated TF footprint remodeling at a developmentally important C4 regulatory locus. Together, these results suggest that attenuation of parental cell-type specificity is concomitant with remodeling of TF occupancy at developmentally relevant regulatory elements in hybrids exhibiting heterosis.

## DISCUSSION

Understanding heterosis requires resolving how divergent parental regulatory programs are combined within the individual cell types that generate hybrid phenotypes. Here, single-cell chromatin accessibility profiling across three maize inbred lines and their six reciprocal F1 hybrids revealed that hybridization does not extensively alter cell-type composition or globally expand the accessible chromatin landscape ^15^. Instead, hybrids showed widespread attenuation of parental cell-type specificity. Regulatory elements that were preferentially accessible in particular parental cell types generally retained a residual bias towards those cellular contexts, but their accessibility became more diffusely distributed across other hybrid cell types. Thus, hybridization primarily broadened the cellular deployment of parental regulatory activity rather than redirecting regulatory elements towards alternative cell identities. This distinction would be obscured in bulk measurements and identifies cell-type specificity as an important dimension of regulatory variation in hybrids.

Our haplotype-resolved analyses further showed that sequence conservation does not necessarily imply conservation of chromatin accessibility. Most ACRs were positionally conserved between parental haplotypes, consistent with a broadly shared regulatory backbone, whereas IS and PAV ACRs were preferentially located in distal and intergenic regions. These structurally divergent non-genic elements were also frequently associated with GWAS and eQTL signals, reinforcing the contribution of distal regulatory variation to phenotypic diversity ^50,61,62^. However, many sequence-conserved ACRs also differed in chromatin accessibility status between parents, indicating that regulatory divergence frequently arises without large structural changes. Such ACRs may instead be shaped by local nucleotide polymorphisms, differential TF recognition or variation in the surrounding *trans*-regulatory environment ^33,63,64^. The enrichment of CPP motifs in accessible-status-variable ACRs, together with their relatively high SNV density, suggests that some conserved regulatory elements occupy sequence contexts that are particularly permissive to evolutionary and developmental modulation. ct-loss ACRs shared many of these features, being concentrated in conserved-variable and distal regulatory regions and enriched for CPP and DOF motifs. These properties define a regulatory compartment that is structurally retained between parents but remains plastic in its regulatory activity and cellular deployment.

The attenuation of chromatin accessibility cell-type specificity observed in hybrids provides a potential regulatory counterpart to genetic complementation. Classical models of heterosis focus on dominance, overdominance and epistasis at the level of alleles and their phenotypic effects ^5–12^. Our results suggest that complementation may also occur through the cellular redistribution of regulatory potential. By weakening parental cell-type bias and broadening the distribution of relative accessibility across cell types, hybrids may allow developmental, metabolic or stress-responsive CREs to operate across a wider range of cellular contexts. Such broadening could increase developmental coordination or buffer cell-type-restricted regulatory limitations inherited from either parent. Furthermore, genes near ct-loss ACRs showed reduced expression specificity in an independent maize transcriptomic dataset ^56^. Nevertheless, reduced cell-type specificity should not be assumed to be inherently advantageous. Some broadening events may be neutral consequences of combining divergent genomes, whereas others may disrupt precise regulatory control. Determining whether attenuation contributes to hybrid performance will therefore require identifying the subset of broadened regulatory elements whose perturbation alters heterotic traits.

Our cell-type-resolved inheritance analyses clarify why chromatin accessibility can appear constant or additive at the tissue level while still exhibiting extensive regulatory complexity. Constant and additive modes were the most frequent inheritance states overall, consistent with recent bulk analyses of maize hybrids ^15,16^. Non-additive inheritance, however, was preferentially associated with highly cell-type-specific ACRs, and individual loci often adopted different inheritance modes across cell types. Increasing cell-type specificity was also associated with greater inheritance-mode entropy, indicating that narrowly deployed regulatory elements are especially sensitive to cellular context when parental genomes are combined. Although the global composition of inheritance modes was highly similar across hybrids sharing a parent, individual loci frequently switched modes in different hybrid combinations. These observations suggest that inheritance is not a fixed property of an ACR. Rather, it emerges from interactions among parental divergence, hybrid genetic background and the regulatory environment of each cell type. Bulk measurements may consequently classify a locus as additive even when it exhibits dominant, transgressive or opposing modes in distinct cellular populations. The *cis*-and *trans*-regulatory analysis provides a mechanistic insight for this context dependence. *Cis*-only effects were maintained across cell types more broadly than *trans*-involved effects, consistent with local sequence differences acting across multiple cellular environments. By contrast, *trans*-only ACRs showed the highest cell-type specificity, indicating that *trans*-regulatory divergence is strongly conditioned by cellular identity. ct-loss ACRs were preferentially associated with same-direction *cis* & *trans* effects and with *trans*-only effects but showed little enrichment for *cis*-only regulation. Attenuation of parental cell-type specificity therefore cannot be explained solely by local allelic variation. Instead, these loci appear particularly responsive to the hybrid trans environment, which may reinforce parental *cis* differences or alter accessibility independently of them.

TF footprint analysis connected this attenuation of parental cell-type specificity to candidate changes in TF occupancy. ct-loss ACRs showed a higher propensity for TF footprint gains than ct-stable ACRs, with DOF and VOZ among the most enriched motif families. Gain-bearing loci were linked to genes involved in development, hormone responses and growth, suggesting that TF occupancy remodeling is not randomly distributed across the genome. A footprint-gaining ACR at the *GLK2* locus provided a representative example of this pattern. Although our data do not establish direct DOF regulation of *GLK2* or a causal contribution to heterosis, this locus is notable given previous evidence linking DOF motifs to bundle-sheath cis-regulatory programs and *GLK2* to bundle-sheath differentiation and C4 photosynthetic specialization ^41,60^, highlighting a candidate site of hybrid-associated TF occupancy remodeling. More broadly, our measurements were restricted to an early seedling stage, and the transcriptional consequences of chromatin accessibility broadening were evaluated using independent tissue-level datasets rather than matched single-cell transcriptomic profiles. Extending these analyses across developmental stages, environmental conditions and genetically diverse hybrids, together with targeted perturbation of candidate ACRs and TFs, will be necessary to establish which regulatory changes contribute directly to hybrid vigor.

Together, our results suggest that hybridization preserves a broadly conserved regulatory backbone while reshaping the cellular deployment of a more plastic subset of regulatory elements. At ct-loss ACRs, parental cell-type biases are attenuated through diffuse chromatin accessibility broadening, and the resulting regulatory states are preferentially associated with context-dependent non-additive inheritance, *trans*-involved divergence and TF occupancy remodeling. By placing cellular context at the center of chromatin accessibility inheritance, this framework explains how the same regulatory element can be integrated differently across cell types and hybrids. Our cell-type-resolved analyses further identifies attenuation of cell-type specificity as a pervasive feature of regulatory remodeling in maize hybrids and a candidate mechanism through which parental regulatory variation may contribute to heterosis.

## METHODS

### Plant materials and growth conditions

Seeds of the inbred maize lines B73, Ki3, and Oh43 were obtained from the U.S. National Plant Germplasm System and germinated in a controlled growth room under a 16 h light/8 h dark photoperiod at approximately 26°C. Self-pollinations and reciprocal crosses were performed. The inbred parents (B73, Ki3, and Oh43) and their F1 hybrids (B73 × Ki3 [B×K], Ki3 × B73 [K×B], B73 × Oh43 [B×O], Oh43 × B73 [O×B], Ki3 × Oh43 [K×O], and Oh43 × Ki3 [O×K]) were subsequently grown under the same conditions. Whole aboveground tissues were collected from seven-day-old seedlings at 10 a.m. and immediately flash-frozen in liquid nitrogen. Two independent biological replicates per genotype were prepared for scifi-ATAC-seq library construction. To evaluate seedling heterosis, plant height and whole-plant fresh weight were measured for ten individual plants per genotype.

### scifi-ATAC-seq library preparation and sequencing

Indexed Tn5 transposase complexes were assembled as previously described ^46^ and used for downstream scifi-ATAC-seq library preparation. For nuclei isolation, a single seven-day-old maize seedling was used per genotype. For the three inbred lines (B73, Ki3, and Oh43), one seedling from each line was pooled prior to nuclei isolation, whereas each reciprocal hybrid was processed individually. Specifically, frozen seedling tissue was briefly ground, and approximately 1 ml of the powdered tissue was transferred into a pre-chilled 2-ml microcentrifuge tube. Ice-cold NIB-cutting buffer (10 mM MES–KOH, pH 5.4; 10 mM NaCl; 250 mM sucrose; 0.1 mM spermine; 0.5 mM spermidine; 1 mM DTT; 1% BSA; 0.5% Triton X-100) was then added. Following buffer addition, the suspension was incubated on ice for 2 min, passed through a 40 μm cell strainer (pluriSelect, #43-10040-40), and centrifuged at 300 rcf for 5 min at 4 °C. The supernatant was carefully removed, and the nuclei pellet was gently resuspended in 500 μl NIB-wash buffer (10 mM MES– KOH, pH 5.4; 10 mM NaCl; 250 mM sucrose; 0.1 mM spermine; 0.5 mM spermidine; 1 mM DTT; 1% BSA). The suspension was passed through a 20 μm cell strainer (pluriSelect, #43-10020-40) and carefully layered onto 1 ml of 35% Percoll buffer (35% Percoll in NIB-wash buffer; Cytiva, #GE17-0891-01) in a 2-ml microcentrifuge tube. Nuclei were centrifuged at 300 rcf for 10 min at 4 °C. Following centrifugation, the supernatant was carefully removed and the nuclei pellet resuspended in 200 μl of TAPS buffer (50 mM TAPS–NaOH, pH 8.0; 25 mM MgCl_2_) containing 0.1% Tween-20 and 0.01% digitonin. The suspension was centrifuged at 300 rcf for 5 min at 4 °C, the supernatant discarded, and the pellet resuspended in 200 μl detergent-free TAPS buffer. A 1 μl of nuclei suspension was diluted 50-fold and stained with DAPI (1 mg/mL) (Sigma-Aldrich, #D9542). Nuclei integrity and concentration were assessed using a hemocytometer under fluorescence microscopy. The nuclei suspension was subsequently adjusted to 4,000/μl based on the measured concentration prior to Tn5 tagmentation. Indexed Tn5 tagmentation was performed using a 96-well plate-based combinatorial indexing scheme with 12 ME-A and 8 ME-B barcodes. For plate layout, nuclei from the pooled inbred lines were distributed across two rows of the 96-well plate, whereas nuclei from each reciprocal hybrid were assigned to a single row (**Fig. 1a**). Each well received 1.5 μl of ME-A complexes and 1.5 μl of ME-B complexes, together with 10 μl of nuclei suspension. Tagmentation was carried out at 37 °C for 1 h and terminated by the addition of 20 μL of quenching buffer (10 mM Tris–HCl, pH 7.8; 20 mM EDTA; 2% BSA). All nuclei were pooled and divided into two 2-ml microcentrifuge tubes. Nuclei were pelleted at 300 rcf for 6 min at 4 °C and resuspended in a total of 200 μl diluted nuclei buffer (DNB; 10x Genomics, #2000207; 100 μl per tube). The suspensions were combined into a single tube, passed through a 20 μm cell strainer (pluriSelect, #43-10020-40), and centrifuged under the same conditions. The supernatant was carefully removed, leaving approximately 8 μl of DNB, which was mixed with 7 μl of ATAC buffer B (10x Genomics, #2000193). From the resulting mixture, a total of 10 μl of nuclei suspension (>1 × 10⁶ nuclei) was used as input for scifi-ATAC-seq library preparation. Libraries were prepared using the Chromium Single Cell ATAC v2 (Next GEM) kit (10x Genomics, #1000390) according to the manufacturer’s instructions. The number of final library amplification cycles was determined by quantitative PCR (qPCR) and set to Cq + 1, with a maximum of nine cycles. Libraries were sequenced on an Illumina NovaSeq 6000 platform in dual-index mode, using 8 and 16 cycles for the i7 and i5 indices, respectively.

### Raw reads processing of scifi-ATAC-seq

Raw BCL files were demultiplexed and converted to FASTQ format using *bcl2fastq* (v2.20.0.422; Illumina) with non-default parameters (--minimum-trimmed-read-length=8 --mask-short-adapter-reads=8). The 16-bp i5 bead barcode was extracted and appended to the read names using the *extract* function in *UMI-tools* (v1.1.6) ^65^ with the parameter --bc-pattern=NNNNNNNNNNNNNNNN. Inline Tn5 barcodes were then demultiplexed and appended to read names using *cutadapt* (v5.0) ^66^ with non-default parameters (-e 0.2 -u 5 -U 5). FASTQ files with modified read names were demultiplexed by well ID (i.e., Tn5 barcode combinations) using a custom script, allowing up to one mismatch per 5-bp half of the 10-bp barcode. Reads were subsequently grouped into one parental pool (B73–Ki3–Oh43; A1–B12) and six F1 hybrid sets: B×K (C1–C12), K×B (D1–D12), B×O (E1–E12), O×B (F1–F12), K×O (G1–G12), and O×K (H1–H12). Processed reads were aligned to the *Zea mays* B73 v5 reference genome (AGPv5) ^47^ using *BWA mem* (v0.7.17) ^67^ with non-default parameters (-M). Notably, scaffolds were excluded from AGPv5, while the mitochondrial (Mt) and plastid (Pt) genomes were incorporated from AGPv4 to improve organellar alignment accuracy. High-quality and properly paired reads were retained using *SAMtools view* (v1.21) ^68^ with non-default parameters (-bhq 10 -f 3). To further ensure mapping specificity, reads carrying the “XA” tag were discarded only if they exhibited low mapping quality (MAPQ < 30) and multiple alternative alignments with comparable edit distances (≤3 mismatches). Additionally, each cell barcode (bam tag ‘BC’) was compared against the 10x Genomics whitelist and retained if it had a Hamming distance of at most two (i.e., ≤2 mismatches). Deduplication was performed per barcode using *picard MarkDuplicates* (v2.18.29; http://broadinstitute.github.io/picard/) with non-default parameters (BARCODE_TAG=BC, MAX_FILE_HANDLES=1000, REMOVE_DUPLICATES=true, ASSUME_SORT_ORDER=coordinate). Finally, BAM alignments were converted to single-base resolution Tn5 insertion sites in BED format by shifting the start coordinates of the forward and reverse strands by +4 and −5, respectively. Only unique insertion sites per cell were retained for downstream analysis.

### Nuclei calling

Nuclei were filtered based on quality metrics using the R package *Socrates* ^32^. To estimate the fraction of reads in peaks (FRiP), the *callACRs* function was applied to bulk Tn5 integration sites with non-default parameters (genomesize=1.6e9, shift=−75, extsize=150, fdr=0.1). The proximity of Tn5 insertions to transcription start sites (TSSs) was quantified using a 2-kb window centered on the TSS via the *buildMetaData* function, generating the corresponding metadata file. Barcodes were identified as putative nuclei if they met the following criteria: (i) ≥1,000 Tn5 insertions per nucleus; (ii) ≥20% of insertions within 2 kb of a TSS, with a standardized proximity greater than two standard deviations below the mean (i.e., Z-score ≥ −2); (iii) FRiP score > 0.2, with a Z-score ≥ −2; and (iv) <5% of insertions mapping to organelle genomes. To further exclude barcodes representing broken nuclei or ambient background, sparse matrices were constructed using the *generateMatrix* function with a 500-bp window size. Spearman’s correlations were then computed between each barcode and two term-frequency inverse-document-frequency (TF-IDF) normalized aggregate profiles: one derived from non-background barcodes (top 1,000 by Tn5 insertions) and another from background barcodes (<100 insertions). Barcodes more strongly correlated with the non-background profile than with the background were retained as high-confidence nuclei for downstream analysis.

### Inter-genomic SNVs calling

To detect sequence variants among B73, Ki3 and Oh43, whole-genome alignments were generated using *AnchorWave* (v1.2.6) ^69^ by aligning the Ki3 and Oh43 genomes separately to the AGPv5 genome ^47^ with the *genoAli* command. The resulting MAF files were converted to GVCF format using the *MAFToGVCF* plugin in *TASSEL* (v5.2.94) ^70^. GVCFs were then consolidated into a GenomicsDB database using *GATK GenomicsDBImport* (v4.3.0.0), and joint genotyping was performed with *GATK GenotypeGVCFs* (v4.3.0.0) using non-default parameters (--stand-call-conf 0 --ploidy 1) ^71^. Indels and non-biallelic sites were excluded, resulting in a final set of 6,362,680 bi-allelic SNVs for downstream analysis.

### Identification of PAV sequences

Presence/absence variation (PAV) sequences among the B73, Ki3, and Oh43 genomes were identified using a sliding-window strategy, as previously described ^72^. For each pairwise comparison, the reference genome was divided into 500-bp overlapping windows with a step size of 100 bp via *bedtools makewindows* (v2.31.1). Each 500-bp window was aligned against both the reference and the compared genome using *BWA mem* (v0.7.17) ^67^ with non-default parameters (-w 500 -M). To define reference-specific sequences, we required that each window could be aligned to the reference genome with 100% primary alignment coverage and MAPQ ≥ 30, but either failed to align to the compared genome or aligned with <25% coverage. Adjacent overlapping windows meeting these criteria were subsequently merged into continuous PAV regions using *bedtools merge* (v2.31.1). This procedure was performed in both directions for each pair of genomes (B73 vs Ki3, B73 vs Oh43, and Ki3 vs Oh43), yielding sets of genome-specific PAV regions.

### Genotype assignment

To genotype single nuclei across both parental and hybrid samples, we applied distinct approaches optimized for each dataset. All analyses were performed on BAM files preprocessed with the *WASP* (v0.3.4) pipeline ^49^ to mitigate reference mapping bias. For the parental pool (B73, Ki3, and Oh43), we applied *Demuxlet* ^73^ under default settings, using the set of SNVs identified in the “inter-genomic SNVs calling” step. Only droplets classified as singlets were retained for downstream analysis. For F1 hybrid samples, although nuclei were initially assigned based on well-specific Tn5 barcodes, a customized allele-specific read count–based framework was applied to eliminate potential mislabeling due to index hopping, inter-sample signal leakage, or sequencing noise. Specifically, for each F1 cross, we selected from the full set of 6,362,680 bi-allelic homozygous SNVs where the two expected parental lines (P1 and P2) carry distinct alleles. These informative SNVs were used to count the number of reads supporting either the P1-or P2-specific allele per barcode. A Bayesian classifier was then applied to assign each nucleus to one of three genotype classes: P1-like, P2-like, or F1 hybrid by modeling allele counts as a binomial distribution. The F1 hybrid class assumed a 1:1 allelic ratio, and empirical priors were set to 0.05, 0.05, and 0.90 to reflect the expected dominance of hybrid nuclei in the dataset. Posterior probabilities were computed via Bayes’ theorem, and barcodes with ≥30 informative reads and ≥0.9 posterior for the hybrid class were retained. To further remove potential inclusion of non-target F1 nuclei, we screened for allele-level evidence of the third, non-contributing inbred using informative SNVs where the two expected parents were homozygous for the same allele and the non-contributing inbred line carried the alternative homozygous allele. For each barcode, we computed the proportion of reads matching the non-contributing allele among total reads covering these informative SNVs. Nuclei with >5% non-contributing allele proportion were excluded from downstream analyses.

### Genotyping quality control

To validate genotype assignments, we compared SNV profiles inferred by the genotype-aggregated counts from scifi-ATAC-seq to the reference genotypes derived from whole-genome alignments described above. BAM files were subsetted using *subset-bam* (v1.0.0; https://github.com/10XGenomics/subset-bam) to retain high-confidence nuclei for each genotype. Variant calling was performed using *freebayes-parallel* (v1.3.8) ^74^ with 24 threads across 5 Mb intervals of the maize AGPv5 genome, using non-default parameters (--strict-vcf -n 4 -q 10 --limit-coverage 10000). Variant calls obtained from scifi-ATAC-seq were merged with those derived from whole-genome alignments using *bcftools merge* (v1.21) ^75^, retaining only ACR-localized SNVs shared between both datasets. A genetic relationship matrix was then computed from the combined variant set using the *snpgdsGRM* function in the *SNPRelate* R package, for filtered biallelic variants (maf=0.05, missing.rate=0; via *snpgdsSelectSNP*) with the correlation-based method (method=“Corr”). In parallel, we evaluated SNV concordance at the single-nucleus level. Allele counts for each nucleus were extracted using *vartrix* (v1.1.22; https://github.com/10XGenomics/vartrix) with non-default parameters (--scoring-method coverage --mapq 30), using the same ACR-localized SNV set as for the genotype relationship matrix. Allele ratios (alt / [ref + alt]) were computed per SNV per cell and discretized into genotype labels: ≤0.2 as 0/0, 0.2–0.8 as 0/1, and ≥0.8 as 1/1. For each nucleus, a similarity score (0–1) was computed by comparing its genotype profile to each parental reference, defined as the proportion of matching genotypes across all valid SNVs (i.e., non-missing in both nucleus and reference). Cells with fewer than 100 valid SNVs were excluded from analysis.

To further independently validate genotype identity, we also assessed chromatin accessibility similarity. Briefly, ACRs were identified from unique Tn5 insertion sites using *MACS2 callpeak* (v2.2.4) ^76^ with non-default parameters (-g 1.6e9 --extsize 150 --shift −75 --nomodel --keep-dup all --qvalue 0.05) and subsequently merged using *bedtools merge* (v2.31.1). Read counts overlapping the merged ACR set were quantified for each genotype replicate using *bedtools multicov* (v2.31.1). The resulting count matrix was scaled using the *cpm* function from the *edgeR* R package (log=TRUE, prior.count=5) ^77^, followed by quantile normalization with *normalizeQuantiles* from the *limma* package. Batch effects across libraries were removed using *removeBatchEffect* (*limma*), and pairwise Spearman’s correlations between genotype replicates were estimated using the base R *cor* function.

### Nuclei clustering

Integrated nuclei clustering across nine genotypes × two biological replicates was performed using *Socrates* R package ^32^. Sparse binary matrices of nuclei × 500-bp genomic bins were first filtered to remove bins accessible in <0.25% or >99.5% of nuclei, as well as nuclei with fewer than 100 accessible features, using the *cleanData* function with non-default parameters (min.c=100 min.t=0.0025 max.t=0.005). The filtered matrix was transformed using TF-IDF algorithm with L2 normalization (*tfidf*, doL2=T). Dimensionality reduction was performed on the top 40% most variable features using a modified *reduceDims* function, with singular vectors discarded due to high correlation with any sequencing-quality metric, including nuclei read depth, transcription start site enrichment, fraction of reads in peaks, or proportion of organellar reads (method=“SVD”, n.pcs=8, num.var=ceiling(nrow(soc.obj$counts)*0.4), cor.max.depth=0.5, cor.max.pTSS=0.5, cor.max.FRiP=0.5, cor.max.pOrg=0.5, scaleVar=T, doL2=T). Pool and genotype effects were removed using the *RunHarmony* function from the *Harmony* R package with non-default parameters (theta=2, sigma=0.1, lambda= 0.1, vars_use=“pool_geno”, nclust=50, max.iter=30, return_object=T). The dimensionality of the nuclei embedding was further reduced with Uniform Manifold Approximation Projection (UMAP) using the *projectUMAP* function with non-default parameters (metric=“cosine”, k.near=30). Clusters of nuclei with similar chromatin accessibility patterns were identified using the *callClusters* of *Socrates* package with non-default parameters (res=1.0, k.near=30, cleanCluster=F, cl.method=4, e.thresh=3, threshold=3, min.reads=1e6, m.clst=50).

### Cell-type annotation

To assign cell types for each cluster, gene activity score was estimated from unique Tn5 integration sites overlapping gene bodies and extending 500-bp upstream of TSSs using the *estGeneActivity* function of *Socrates* and normalized to a total of 10,000 per nucleus. Cell types were initially assigned to clusters based on the evaluation of known marker gene activity on UMAP projections and the recovery of known cell-type markers among *de novo* identified markers. Known cell-type marker genes were primarily curated from a previous maize scATAC-seq study ^32^ and supplemented with additional markers from other literature (**Supplementary Table 4**). To account for cell-cycle-associated effects on clustering and cell-type annotation, we compiled a set of 328 (**Supplementary Table 5**) previously characterized maize cell-cycle-phase-specific genes^78^. For each cell, phase scores were derived as the mean gene activity of the corresponding marker set. Cells with zero scores across all phases were defined as G0. To evaluate the significance of phase scores, permutation-based normalization was performed. For each phase, gene sets of matched size were randomly sampled from the pooled cell cycle markers to generate empirical null distributions over 1,000 permutations. Phase-specific *Z*-scores were calculated by standardizing the observed mean gene activity relative to the corresponding permutation-derived distributions. Cells not classified as G0 were assigned to the cell cycle phase corresponding to the highest phase-specific *Z*-score. Cell-type annotations were finalized by integrating predicted cell-type identities with inferred cell cycle stage information. Cell-type composition was calculated as the proportion of nuclei assigned to each annotated cell type within each genotype-replicate sample. For each cell type, hybrid–inbred differences in cell-type proportion were estimated using a linear model with group (inbred or hybrid) as the main effect and biological replicate as a covariate. Equivalence test was performed on the estimated group effect using two one-sided tests (TOST) ^79^ with a predefined margin of ±0.05 in cell-type proportion. *P* values were adjusted across cell types using the Benjamini–Hochberg method. To assess whether gene ontology (GO) enrichment profiles were consistent with expected cell-type functions, gene set enrichment analysis (GSEA) was performed using the R package *fgsea* ^80^, following a previously described framework ^39^. For each cell type, a balanced reference panel was constructed by uniformly sampling nuclei from each cluster, with the total number of reference nuclei set to the mean cluster size. Gene-level accessibility profiles were aggregated across nuclei within each cell type and within the reference panel to generate pseudo-bulk profiles. Differential accessibility between each cell type and the reference panel was assessed using *edgeR* R package ^77^. For each cell type, genes were ranked in decreasing order of log_2_ fold change relative to the reference panel and used as input for gene set enrichment analysis. GO terms with fewer than 10 or more than 600 genes were excluded. Enrichment significance was evaluated using 10,000 permutations, and gene sets with FDR < 0.05 were considered significant (**Supplementary Table 6**).

### Genotype-aware realignment

To accurately capture genotype-specific chromatin accessibility, sequencing reads from each genotype × cell type × replicate combination were re-aligned to the corresponding parental reference genome (B73, Ki3, or Oh43) ^47^ for inbred samples, or to a synthetic diploid assembly formed by concatenating the two parental assemblies for hybrids. In the hybrid reference, chromosome names were prefixed with haplotype-specific tags (e.g., B73_chr1, Ki3_chr1 or Oh43_chr1) to distinguish parental haplotypes. For inbreds, read alignment and subsequent BAM processing were carried out exactly as described above in the “scifi-ATAC-seq raw data processing” section. For hybrids, BAM filtering differed from the parental procedure. Specifically, properly paired reads (-f 3) were streamed from *SAMtools view* (v1.21) ^68^ and examined for mapping quality and alternative alignments recorded in the “XA” tag. Alignments were retained based on the following criteria: (i) no alternative alignments or no cross-parent alternatives: discard if MAPQ < 10; keep if MAPQ > 30; otherwise (10 ≤ MAPQ ≤ 30) keep only when ≤1 same-parent alternative had a comparable edit distance (≤3 mismatches); (ii) with cross-parent alternatives: keep if MAPQ > 30; otherwise keep only when MAPQ ≤ 30 with exactly one alternative mapping to the same chromosome of the other parent. The remaining steps, including cell barcode correction, PCR duplicate removal, and conversion to single-base Tn5 insertion sites were performed exactly as described above.

### ACR identification

Using aggregated Tn5 insertion counts from the re-alignment, ACRs were called using *MACS2 callpeak* (v2.2.4) ^76^ separately for each combination of cell type, genotype, and biological replicate, with non-default parameters (-g 1.6e9 --nomodel --keep-dup all --extsize 150 --shift −75 --qvalue 0.05) for parental inbreds and -g 3.2e9 for hybrids. We then stringently filtered candidate ACRs to minimize false positive rate by retaining only those that were accessible in a significant fraction of the cells compared to background regions. To achieve this, we randomly selected a control set of genomic regions equal in number and size to the candidate ACRs from mappable genomic regions with *bedtools shuffle* (v2.31.1), which were defined as regions covered by at least one read in a simulated dataset generated via *wgsim* (v1.21; https://github.com/lh3/wgsim) with non-default parameters (-N 400000000 -1 150 -2 150 -d 300). For each control region, we calculated the fraction of cells with at least one Tn5 insertion using *bedmap* (v2.4.41) ^81^ with non-default parameters (--echo-map-id-uniq) and used these fractions to construct an empirical cumulative distribution function (ECDF) as the null model. For each candidate ACR, we then quantified the fraction of accessible cells and assessed its significance relative to this ECDF. ACRs with FDR < 0.01 that were accessible in at least 1% of cells (minimum one cell) and reproducibly detected in both biological replicates were retained for downstream analysis. To compile a non-overlapping union set of fixed-width (501 bp) ACRs for each genotype or haplotype, we combined ACRs from all cell type × replicate combinations and extended the ACR summits by 250 bp on either side. Overlapping ACRs were resolved through an iterative merging and removal procedure, as previously described ^33,82^. Briefly, the concatenated ACR sets were ranked by their statistical significance (based on −log_10_ q-value), and the most significant ACR was retained while any directly overlapping ACRs were progressively removed until no further overlaps remained. For hybrid samples, this procedure was applied separately to each parental haplotype.

### Downsampling analysis of global ACR properties

To compare global ACR properties between inbred and hybrid genotypes, all pseudo-bulk ATAC-seq reads were aligned to maize AGPv5 genome and filtered with *WASP* (v0.3.4) ^49^ to reduce reference-mapping bias. Tn5 insertion sites were pooled across cell types within each genotype, and the minimum Tn5 insertion count across genotypes was used as the shared depth. Each genotype was downsampled to 10%, 25%, 50% and 100% of this shared depth using reservoir sampling with five random seeds. ACRs were called from each downsampled Tn5 BED file using *MACS2 callpeak* (v2.2.4) ^76^ with -g 1.6e9 --nomodel --keep-dup all --extsize 150 --shift −75 --qvalue 0.05. ACR number and length were then quantified for each downsampling replicate.

### ACR classification

To enable systematic cross-genome comparison of ACRs across maize genotypes, we first constructed pairwise whole-genome chain files. Each genome pair was aligned using *minimap2* (v2.30-r1287) ^83^ with non-default parameters (-cx asm5 --cs), and the resulting pairwise PAF alignments files were converted into chain files using *transanno* (v0.4.5; https://github.com/informationsea/transanno). Starting from a reference-centric ACR set, all ACRs were represented by their summit positions. Summit coordinates were transferred to the target genome using *liftOver* (v482) ^84^. ACRs whose summits could be successfully transferred were classified as directly-mapped ACRs and considered sequence-conserved. Unmapped summits overlapping reference-specific PAV intervals were classified as PAV ACRs, reflecting loci present in the reference genome but absent from the target genome. The remaining unmapped summits were further evaluated using a sequence-based rescue strategy to identify structurally shifted ACRs. Specifically, each summit was expanded to a 500-bp window, from which 150-bp flanking edge sequences were extracted and independently aligned to the target genome using *minimap2* (v2.30-r1287) ^83^ with non-default parameters (-x sr -k 15 -w 5 --secondary=no). Only high-confidence alignments (MAPQ ≥ 30 and query coverage ≥ 70%) were retained. The relative genomic positions of the mapped left and right edges were then integrated to infer local structural relationships between genomes. Based on the deviation between the observed and expected inner distances of the mapped flanking edges (expected inner distance = 200 bp), ACRs were classified into four categories. ACRs with minimal deviation (≤10 bp) were classified as minor-shifted. ACRs showing moderate deviations (10 bp < deviation ≤ 2 kb) were classified as small-insertion, whereas those with larger but bounded deviations (2 kb < deviation ≤ 50 kb) were classified as large-insertion. Loci exhibiting excessive deviations (>50 kb), discordant or cross-chromosomal mappings, or incomplete flank recovery were classified as ambiguous. Minor-shifted ACRs were merged with directly-mapped ACRs to define the conserved ACR set, representing loci with preserved genomic positioning across genomes. For downstream analysis, small-and large-insertion ACRs were combined and collectively referred to as insertion-shifted (IS) ACRs, whereas PAV ACRs were retained as a separate category. Ambiguous ACRs were excluded from downstream quantitative analyses. To assess conservation at the accessibility level, conserved, and IS ACRs were merged per haplotype and compared reciprocally between genomes. Using coordinate overlaps in the corresponding genomes, defined as the presence of any genomic overlap (≥1 bp) between ACR intervals, ACRs were classified as shared if an overlapping ACR was detected in the other genome, or as variable if no overlap was observed. ACRs were also categorized with genomic context using haplotype-matched gene models (longest transcript per gene). For each ACR, distance to the nearest gene and base-pair overlap with gene bodies, exons and introns were computed. ACRs were classified as intergenic ACRs if no gene overlap was detected and the nearest gene was > 2 kb away, or proximal ACRs otherwise. For genic ACRs, loci with gene overlap < 50% of ACR length were labeled proximal ACRs, and remaining ACRs overlapping a TSS were labeled as TSS ACRs, whereas exonic ACRs or intronic ACRs status was assigned if the corresponding feature covered > 50% of ACR length.

To identify ctACRs, we constructed a non-redundant union non-PAV ACR set across two parental haplotypes following the iterative merging and removal procedure described in the “ACR identification” section. Chromatin accessibility of union non-PAV ACRs and haplotype-specific PAV ACRs was quantified across all parental and hybrid samples by counting reads overlapping each ACR using *bedtools multicov* (v2.31.1). For hybrid samples, reads overlapping non-PAV ACRs were assigned to each parental haplotype using haplotype-tagged pseudo-F1 reference coordinates and summed to obtain total hybrid accessibility, whereas PAV ACRs were quantified from their corresponding haplotype only. Raw counts were normalized to counts per million (CPM) using sample-specific library sizes computed from the total accessibility across all ACRs, including both non-PAV and PAV loci. Cell-type specificity of each ACR was estimated via the

Tau metric ^33,85^, calculated from CPM-normalized accessibility values across cell types. ctACR was defined if it satisfied both mean Tau ≥ 0.7 and ΔTau ≤ 0.15, where ΔTau was defined as the difference in Tau values across biological replicates for the same ACR. ACRs with mean Tau < 0.7 were classified as nctACR, and remaining loci were labeled as unresolved ACR.

### Motif analysis

Motif enrichment analysis was performed across ACR categories. For each category, foreground (FG) regions were defined as ACRs belonging to the focal category, whereas background (BG) regions comprised all remaining ACRs. To minimize gene-body sequence bias, motif enrichment analyses were restricted to intergenic, proximal, and TSS ACRs. ACR sequences were extracted from the haplotype-matched reference genome, and motif scanning was performed using *FIMO* from the *MEME* suite (v5.5.5) ^86^ with the non-redundant core plant motif database in JASPAR 2024, applying a significance threshold of *P* < 1 × 10^-4^. Motif enrichment was assessed by comparing the number of FG and BG ACRs containing each motif using two-sided Fisher’s exact test. Multiple testing correction was performed using the Benjamini–Hochberg procedure, and motifs with FDR < 0.05 were considered significantly enriched or depleted. To assess motif-associated sequence polymorphism, SNV density was quantified within motif bodies in sequence-conserved ACRs, thereby minimizing confounding from large-scale structural variations. Overlapping *FIMO* hits assigned to the same motif ID were merged to avoid double-counting. For each motif interval, SNV density was calculated as SNV count divided by motif length, and local background SNV density was calculated from the remainder of the corresponding ACR after excluding the motif interval. Background-corrected SNV density was defined as motif-body SNV density minus local background SNV density. To assess whether differences in polymorphism could be explained by motif sequence composition, trinucleotide-specific SNV probabilities were estimated from non-motif positions within sequence-conserved ACRs. The expected SNV density for each motif was then calculated from the trinucleotide contexts of its genomic motif positions, and sequence-context-corrected SNV density was defined as observed minus expected density. Motifs represented by fewer than 50 instances were excluded from ranking. To predict the effects of CPP-site variants on motif binding, SNVs overlapping *FIMO*-defined SOL1, TSO1, TCX6, TCX2 and TCX3 motif instances were evaluated using *PERFECTOS-APE* ^54^. JASPAR motif matrices were used to score the B73 reference and alternative alleles for each variant–motif pair, and effect magnitude was summarized as the absolute log_2_ ratio of reference-to alternative-allele motif P values. Pairs were classified as having strong predicted effects when at least one allele had a motif *P* < 5 × 10^-4^ and the allele-specific *P* values differed by at least fivefold.

### Hybridization-associated attenuation analysis

To assess whether differences in sequencing depth contributed to the observed reduction in cell type specificity in hybrids, inbred and hybrid datasets were downsampled to matched read depths before recalculating CPM and Tau values. To examine how attenuated ACRs were distributed across hybrid cell types, biological replicates were first averaged for each genotype–cell type combination. For each ACR and genotype, accessibility proportion in each cell type was calculated as the CPM value in that cell type divided by the summed CPM across all cell types. Parental accessibility proportion was calculated as the mean value of the two parental inbreds, and hybrid accessibility proportion was calculated as the mean value of the two reciprocal hybrids. The parental source cell type of each ACR was defined as the cell type with the highest mean parental accessibility proportion. To exclude ACRs with ambiguous source assignment, we required a difference of at least 0.05 between the highest and second-highest parental accessibility proportions. For each parental source cell type, hybrid accessibility profiles were summarized by averaging hybrid accessibility proportions across ct-loss ACRs assigned to that source cell type. Changes in accessibility proportion were calculated as hybrid mean proportion minus parental mean proportion.

To quantify changes in the cellular accessibility breadth of ct-loss ACRs, replicate-level raw fragment counts across the four genotypes (two parents and two reciprocal hybrids) and 14 cell types within each cross were jointly fitted with a two-component negative-binomial mixture model:

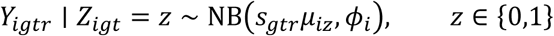

where *Y*_*igtr*_ is the fragment count for ACR *i*, genotype *g*, cell type *t* and replicate *r*, and *Z*_*igt*_ is the corresponding latent accessibility state shared across replicates. *s*_*gtr*_ is the sample-specific size factor, *µ*_*iz*_ is the ACR-specific mean for state *z*, and *ϕ*_*i*_ is the ACR-specific dispersion; size factors and dispersions were estimated using *DESeq2* ^87^. The higher-mean component was designated the accessible state, and genotype–cell-type contexts with *P*(*Z*_*igt*_ = 1) ≥ 0.5 were classified as accessible. ACRs with fewer than 20 total fragments across all samples within a cross or unsuccessful model fits were excluded. Cellular breadth was defined as the number of accessible cell types per genotype and averaged across the two inbreds or corresponding reciprocal hybrids.

### Differential accessibility and inheritance mode analysis

To focus specifically on putative *cis*-regulatory elements, exonic ACRs were excluded from all downstream analysis, and only non-PAV ACRs shared between parental genomes were retained for differential accessibility and inheritance-mode analysis. DARs were identified separately for each cell type from raw ACR read counts using *DESeq2* ^87^. ACRs with total read counts ≥ 4 in each contrast were tested. Reciprocal hybrids (e.g., B×K vs K×B) were first compared to assess potential parent-of-origin effects. ACRs were considered to exhibit reciprocal effects only if they satisfied FDR < 0.05 and |log_2_ fold change| ≥ 1. No genome-wide reciprocal effects meeting these criteria were detected; therefore, reciprocal hybrids were pooled and treated as a single hybrid group for subsequent analysis. Differential testing was then performed for the parental comparison (e.g., B73 vs Ki3) and for each parent versus the pooled hybrid (e.g., B73 vs hybrid and Ki3 vs hybrid). For the parental contrast, SPAs were defined using CPM thresholds (mean CPM ≥ 1 in one parent and < 0.1 in the other) and remaining significant ACRs (FDR < 0.05) were stratified by fold-change magnitude (2–4 fold and >4 fold). Parental DARs were summarized per cell type by direction and effect size, and cross–cell type directional consistency was evaluated across all cell types, excluding cell types with fewer than 10 detected parental DARs to ensure robust estimation. Inheritance modes were classified using normalized accessibility values for each parent and the pooled hybrid. Constant ACRs were defined as those for which all pairwise *DESeq2* ^87^ contrasts across all four genotypes (the two parents and two reciprocal hybrids) showed |log_2_ fold change| < 1 and FDR > 0.05. Non-constant ACRs were then hierarchically categorized using a 1.5-fold threshold into overdominant (hybrid > 1.5 × high parent), underdominant (hybrid < low parent / 1.5), additive (hybrid within ±1.5-fold of the mid-parent value), or parental dominance based on proximity to either parent.

To quantify variability of inheritance mode, Shannon entropy was calculated for each ACR. Inheritance modes across all cell types were converted to relative frequencies, and entropy was computed as:

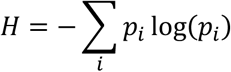

where *p*_*i*_ represents the proportion of cell types assigned to inheritance mode *i*. Entropy values close to zero indicate uniform inheritance classification across cell types, whereas higher values reflect increased mode heterogeneity. Inheritance modes were also compared across different hybrid combinations at the locus level, including B73–Ki3 vs B73–Oh43 and B73–Ki3 vs Ki3–Oh43 hybrids. For each comparison, ACRs were projected onto the coordinate system of the shared parent genome and merged into loci using a 1 bp overlap criterion with *bedtools merge* (v2.31.1). Thus, B73–Ki3 vs B73–Oh43 comparisons were performed in B73 coordinates, whereas B73–Ki3 vs Ki3–Oh43 comparisons were performed in Ki3 coordinates. Inheritance modes were recoded relative to the shared parent (P1), with dominance of the shared parent classified as P1-dominant and dominance of the alternative parent as P2-dominant. For each locus and cell type, the predominant inheritance mode was assigned based on the most frequent ACR-level mode, while loci with ties were labeled as mixed and excluded from strict comparisons. Mode concordance and transition rate were then quantified across loci shared between hybrids.

### *Cis* and *trans* regulatory inference for ACRs

To classify regulatory divergence for each ACR, we jointly modelled parental accessibility and hybrid allelic accessibility within each cell type, as previously described ^56,58,59^. Before model fitting, low-information ACRs were filtered to retain regions with sufficient parental and hybrid allelic read support. Specifically, we required a minimum total count of 10 across the two parents, a minimum total count of 20 across all hybrid samples (including four reciprocal hybrids), at least 10 reads for a hybrid replicate to be considered valid, and support from at least two valid hybrid replicates per ACR. For a given ACR, let *x*_*i*_ denote the accessibility count in the *i*-th P1 replicate (e.g., B73), *y*_*i*_ denote the accessibility count in the *i*-th P2 replicate (e.g., Ki3), *n*_*j*_ denote the total number of allele-informative reads in the *j*-th hybrid replicate, and *z*_*j*_ denote the number of allele-informative reads assigned to the P1 haplotype in the *j*-th hybrid replicate. We assumed:

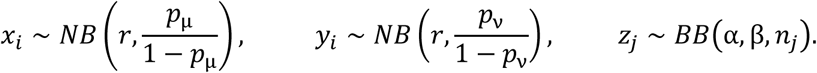

where the marginal distributions of *x*_*i*_ and *y*_*i*_ were assumed to be negative binomial, whereas the marginal distribution of *z*_*j*_ was assumed to be beta-binomial. Additionally, *r* reflects overdispersion relative to a Poisson distribution and was estimated a priori following the previous approach ^88^. For each ACR, four models were fitted by maximum likelihood, and the best-fitting model was selected using the Bayesian information criterion (BIC):

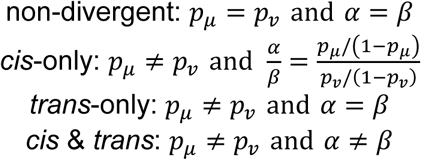

ACRs assigned to the *cis* & *trans* class were further subdivided into four subclasses. We defined *m* as the mean log_2_ allelic ratio for the F1 samples and *n* as the mean log_2_ fold change for the parental values. For the parental data, accessibility counts were first normalized by CPM to account for library-size differences. Opposite-direction *cis* & *trans* effects were considered compensating configurations, whereas same-direction effects were considered reinforcing configurations. The four subclasses were defined as follows:

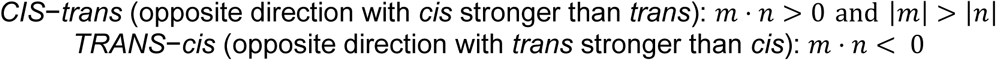

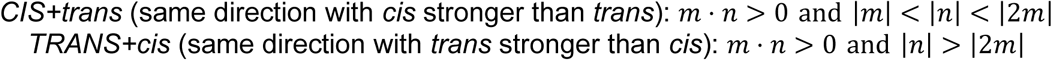

### Cell-type-resolved TF footprinting

TF footprinting was performed using pre-generated cell-type pseudo-bulk BAM files. Parental BAM files had been aligned to their respective NAM-5.0 genome assemblies, whereas F1 BAM files had been aligned to concatenated pseudo-diploid references comprising the two corresponding parental assemblies, with chromosome labelled by parental origin. For each F1 pseudo-bulk, the existing BAM file was partitioned by chromosome prefix into the two parental alleles using *pysam* (v0.23.3), retaining primary mapped alignments with mapping quality ≥10 and excluding secondary and supplementary alignments. Thus, for each cell type, footprint comparisons comprised the two parental samples in their respective genomic backgrounds (P1 and P2) and the corresponding two alleles within the shared F1 nucleus (F1A and F1B). Reciprocal F1s were analyzed independently as replicates of the same parental allele comparison. PCC, MSC2 and PDC were excluded from footprint analyses because only a small fraction of ACRs in these cell types met the coverage requirements described below. Footprint analysis was restricted to cross-specific ACRs paired between the two parental coordinate systems, excluding PAV regions, yielding 64,371 paired ACRs for B73–Ki3 and 67,783 for B73–Oh43. Each ACR was re-centered and expanded to a 2-kb window independently in each parental genome, and the corresponding peak-pairing table was retained for cross-genome comparisons. The 926 JASPAR CORE plant motifs were clustered within TF families using *motifStack* (v1.46.0) ^89^, yielding 227 non-redundant motif signatures. Clustering was restricted to within-family comparisons to avoid collapsing structurally similar motifs from distinct TF families. Motif signatures were scanned independently across each parental 2-kb sequence using *MOODS* (v1.9.4.1) ^90^, assuming a uniform nucleotide background and a significance threshold of *P* = 5 × 10^-5^, with both strands considered. Because scanning was performed on the corresponding parental sequence, motif occurrences were allowed to differ between haplotypes.

For footprint-score calculation, properly paired, primary, same-contig read pairs from each pseudo-bulk were converted to fragments and assigned to a single barcode, such that each pseudo-bulk was treated as one pseudo-cell. Fragment files were imported into *scPrinter* (v1.2.1; *SnapATAC2* v2.8.0 import backend) ^91,92^. Sequence-intrinsic Tn5 insertion bias was modelled separately for B73, Ki3 and Oh43 using the pre-trained *scPrinter* Tn5 bias model applied to genomic sequence alone. *scPrinter* was then used to calculate TF-binding and nucleosome scores (context radius, 100 bp; downsampling factor, 5) and multi-scale footprint scores (99 scales, 2–100 bp). Footprint scores were represented as −log_10_ P, with higher values indicating stronger evidence of local protection from Tn5 insertion. Because footprint scores depend strongly on local sequencing depth, we applied library-specific coverage filtering followed by coverage-stratified normalization. For each parental genome, 10,000 random 2-kb windows were sampled from chromosomes 1–10 after excluding windows overlapping ACRs. For each library, the 95th percentile of fragment coverage across the corresponding background windows was used as the coverage threshold. A region–library pair passed only if coverage exceeded this threshold and contained at least 50 fragments. For each cell type, a paired ACR was retained when at least two of the four conditions (P1, P2, F1A and F1B) passed this filter. Because ≤5.9% of paired peaks met this criterion in PCC, MSC2 and PDC, these cell types were excluded from downstream footprint analyses in both crosses. To further control for coverage-dependent variation, footprint scores were normalized against a coverage-matched empirical null separately for each pseudo-bulk. Within regions containing at least one fragment, the 20% of *scPrinter* TF-binding-score tiles with the lowest scores were treated as putatively unbound background positions. These positions were pooled across regions and divided into 20 quantile bins according to regional fragment coverage. Within each bin, the null distribution was summarized by its median and a robust estimate of the standard deviation, calculated as IQR / 1.349. Each motif occurrence was assigned to the coverage bin of its corresponding region. Each motif occurrence was assigned to the coverage bin of its corresponding region. The motif-level footprint score was calculated by averaging *scPrinter* footprint scores across scales of 2–20 bp within ±5 bp of the motif center and was normalized as:

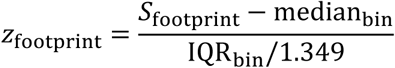

where *S*_footprint_ denotes the average footprint score. Motif occurrences with *z*_footprint_ > 2 were considered footprint positive. This coverage-stratified normalization reduced coverage-dependent variation among pseudo-bulks and enabled comparison of footprint strength across parental and hybrid alleles.

Parent-to-hybrid footprint changes were calculated separately for the two parental alleles (F1A − P1 and F1B − P2). To compare ct-stable and ct-loss ACRs, an ACR–motif pair was classified as gaining at a given score-change threshold when either parental lineage showed gains in at least two cell types, with at least two cell types available for comparison. The fraction of gaining motifs was calculated for each ACR across thresholds of 1–10, and ct-stable and ct-loss ACRs were compared using a one-sided permutation test based on the area under the threshold-response curve. For subsequent ct-loss analyses, high-confidence footprint gains were defined by a score increase ≥2, valid measurements in at least six cell types and gains in >50% of valid cell types in both parental lineages. Motif-family gain enrichment was assessed by Fisher’s exact test followed by Benjamini–Hochberg correction (FDR < 0.05), requiring at least 20 testable ACRs per family. Genes nearest to high-confidence gain-bearing ct-loss ACRs were used for GO enrichment with AgriGO_v2 ^93^ under default settings.

## DATA AVAILABILITY

Raw and processed data generated by this study will be publicly available at National Center for Biotechnology Information Gene Omnibus (NCBI GEO) following peer review.

## CODE AVAILABILITY

The code used for the analyses throughout the manuscript is available via GitHub at https://github.com/lgjiang1/maize_single-cell_heterosis.

## ACKNOWLEDGEMENTS

We wish to thank the Advanced Research Computing at the University of Michigan, Ann Arbor for providing computational resources and technical support. This work was supported by the National Institutes of Health (R00GM144742) and start-up funds from the University of Michigan to A.P.M.

## CONTRIBUTIONS

A.P.M. and L.J. designed and conceived experiments and managed the project. L.J., F.G.C. and J.L. performed the experiments. L.J., F.G.C., M.A.A.M. and A.P.M. analyzed the data. L.J. and A.P.M. wrote the manuscript. All authors approved the final manuscript.

## DECLARATION OF INTERESTS

The authors declare no competing interests.

---

**Supplementary Fig. 1.**
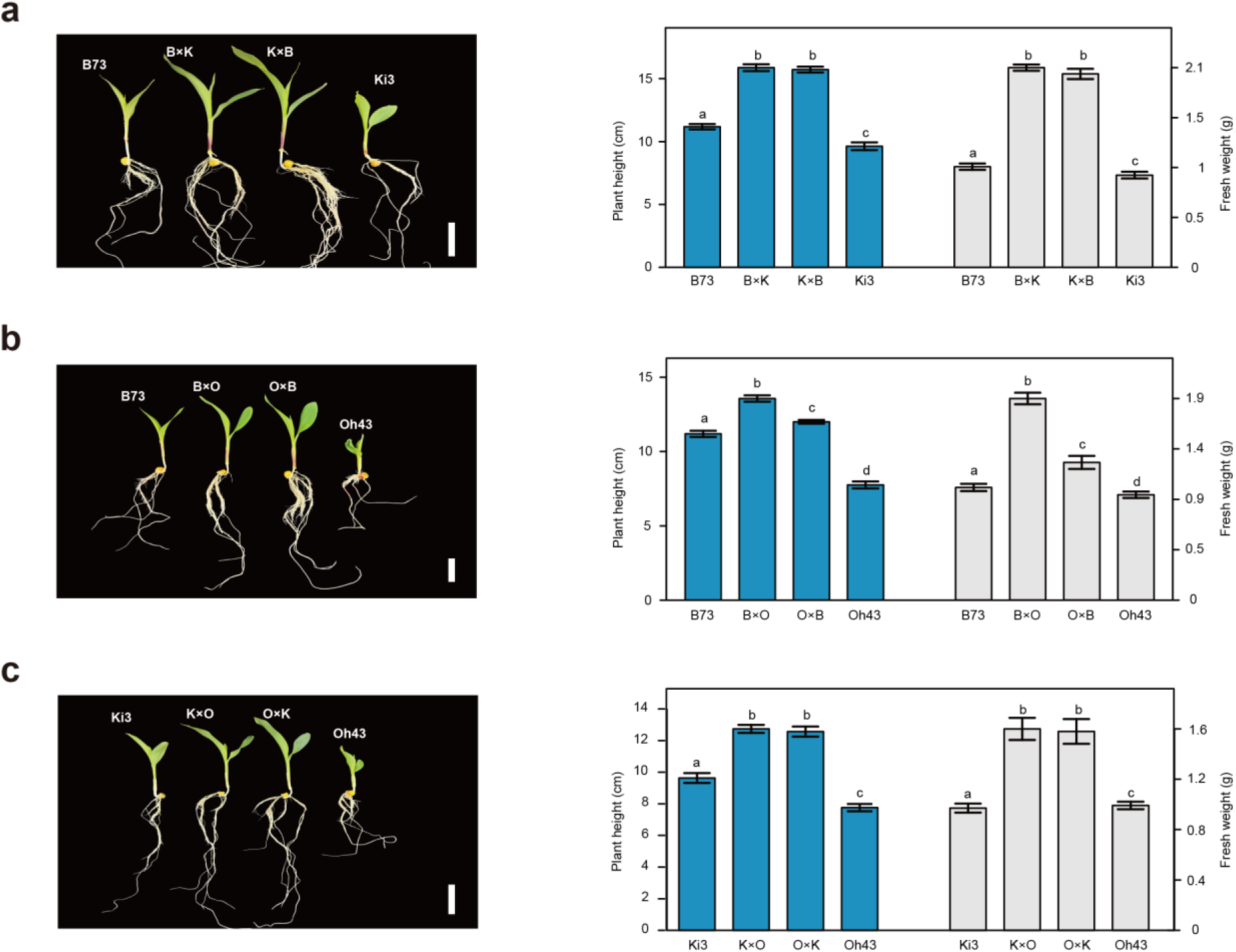
Hybrid vigor in maize seedlings. **a–c**, (Left) Representative images of maize seedlings for parental lines and their reciprocal hybrids. (**a**) B73, B×K, K×B and Ki3; (**b**) B73, B×O, O×B and Oh43; (**c**) Ki3, K×O, O×K and Oh43. Scale bars, 5 cm. (Right) Quantitative analysis of plant height (blue bars) and fresh weight (white bars) for each genotype. Letters denote significant differences among genotypes (Student’s t-test, *P* < 0.05). Error bars represent standard errors of the mean.

**Supplementary Fig. 2.**
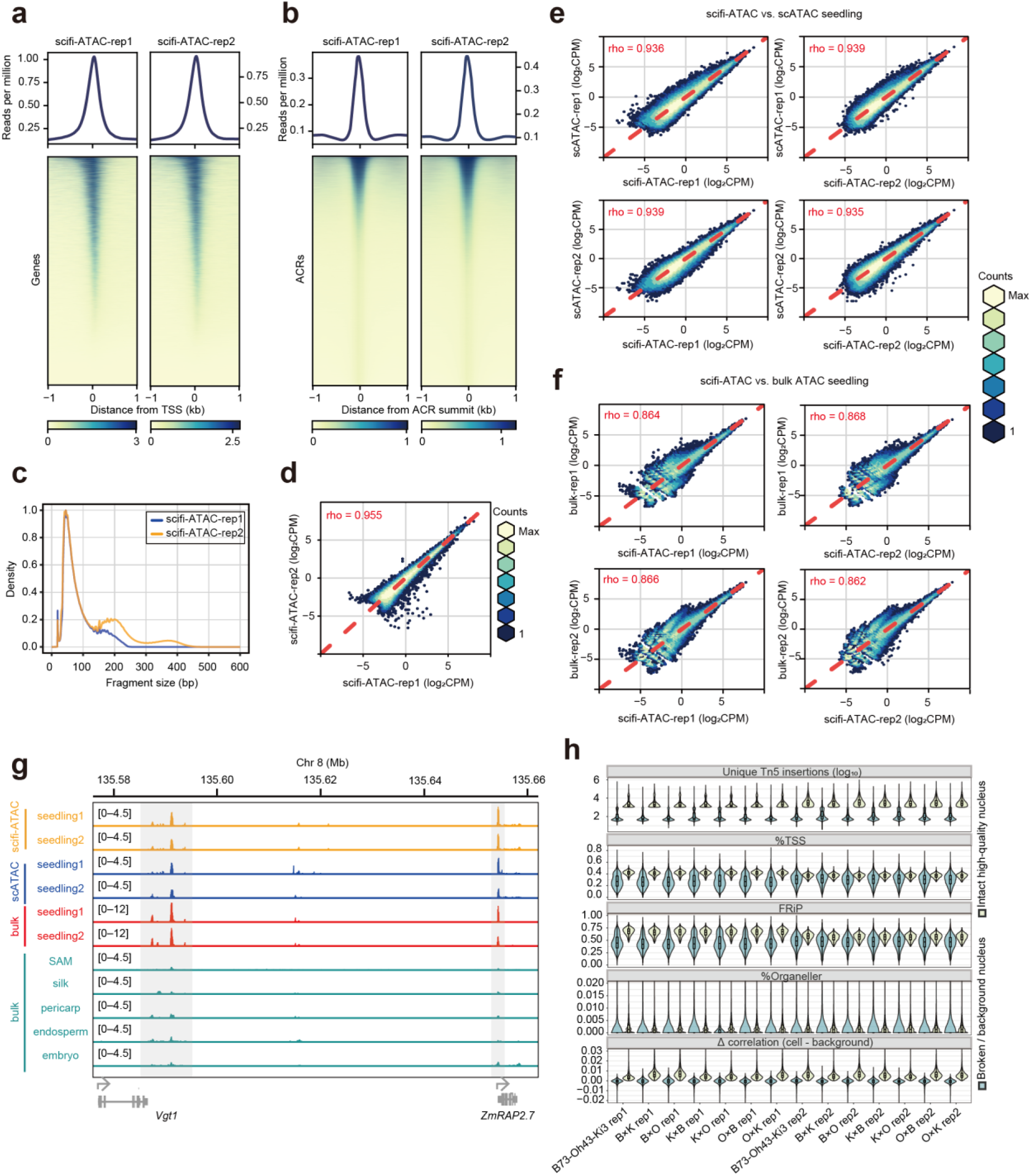
Assessment and quality control of scifi-ATAC-seq data. **a–b**, Metaplot and heatmap of ATAC-seq signals surrounding transcription start sites (TSS, ±1 kb; A) of annotated genes and ACR summits (±1 kb; B) across two scifi-ATAC-seq biological replicates. **c**, Fragment size distributions for each library. **d**, Comparison between two scifi-ATAC-seq biological replicates using log_2_-transformed CPM values across the union of ACRs. Color indicates local point density. **e–f**, Correlation analysis of log_2_-transformed CPM values across the union of ACRs between scifi-ATAC-seq libraries and previously published scATAC-seq (**e**) and bulk ATAC-seq (**f**) seedling datasets. Color indicates local point density. **g**, Genome browser tracks showing the *ZmRAP2.7* gene and its distal regulatory region *Vgt1* (∼70 kb upstream), from scifi-ATAC-seq, scATAC-seq and bulk ATAC-seq experiments. **h**, Barcode quality control filtering metrics.

**Supplementary Fig. 3.**
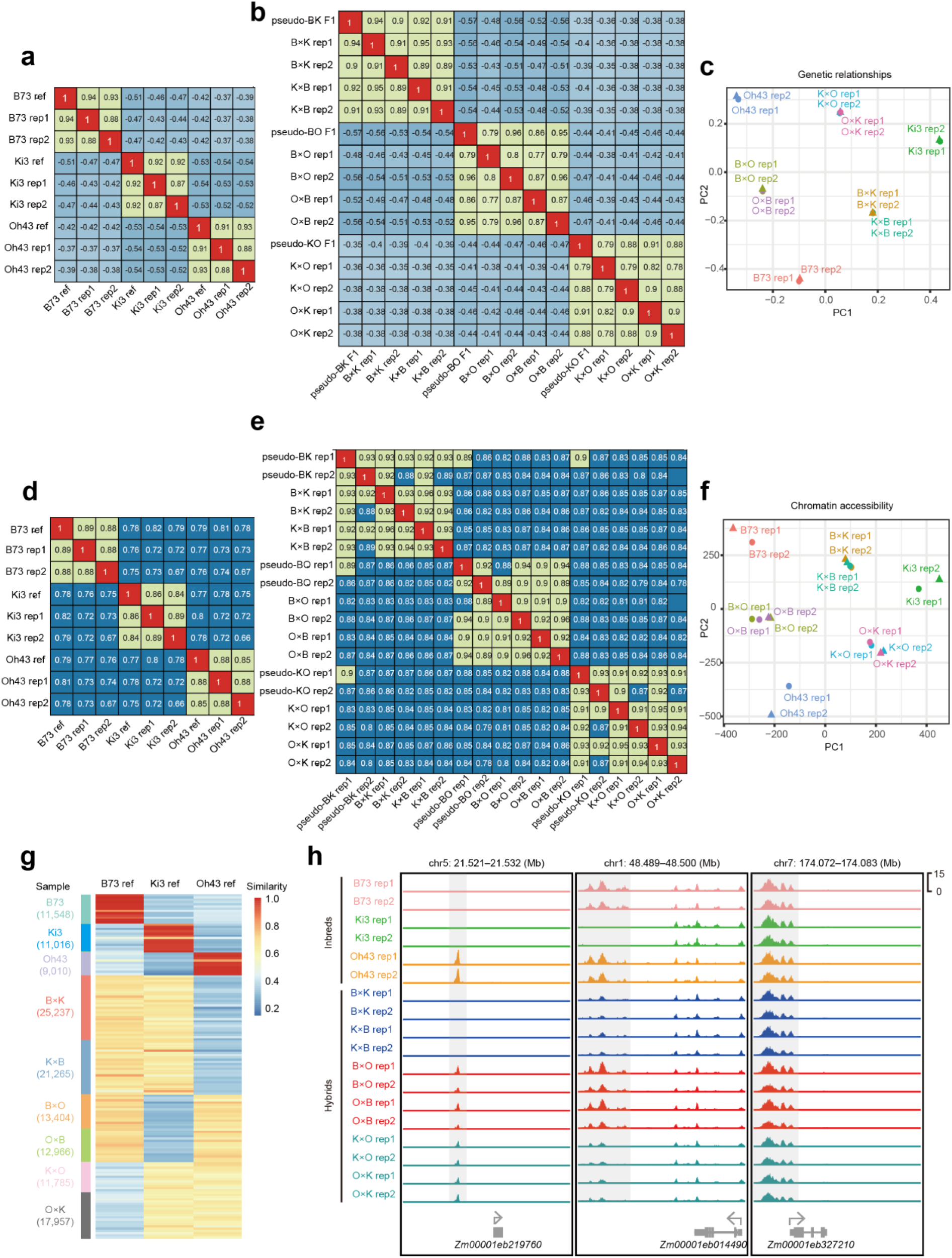
Genotyping quality metrics for scifi-ATAC-seq. **a**, Genetic relationship matrix between scifi-ATAC-seq variants and whole-genome alignments variants for parental inbreds. **b**, Genetic relationship matrix between scifi-ATAC-seq variants and pseudo-F1 variants for hybrids. **c**, Scatter plot of the first two PCs based on genetic relationship. **d**, Comparison of chromatin accessibility profiles between inbred scifi-ATAC-seq and published scATAC-seq profiles of the same genotypes. **e**, Comparison of chromatin accessibility profiles between hybrid scifi-ATAC-seq and pseudo-F1. **f**, Scatter plot of the first two PCs based on chromatin accessibility. **g**, Cell-wise similarity (proportion of matching SNVs) to each parental reference genome. **h**, Representative loci illustrating accurate genotype assignment and expected peak inheritance.

**Supplementary Fig. 4.**
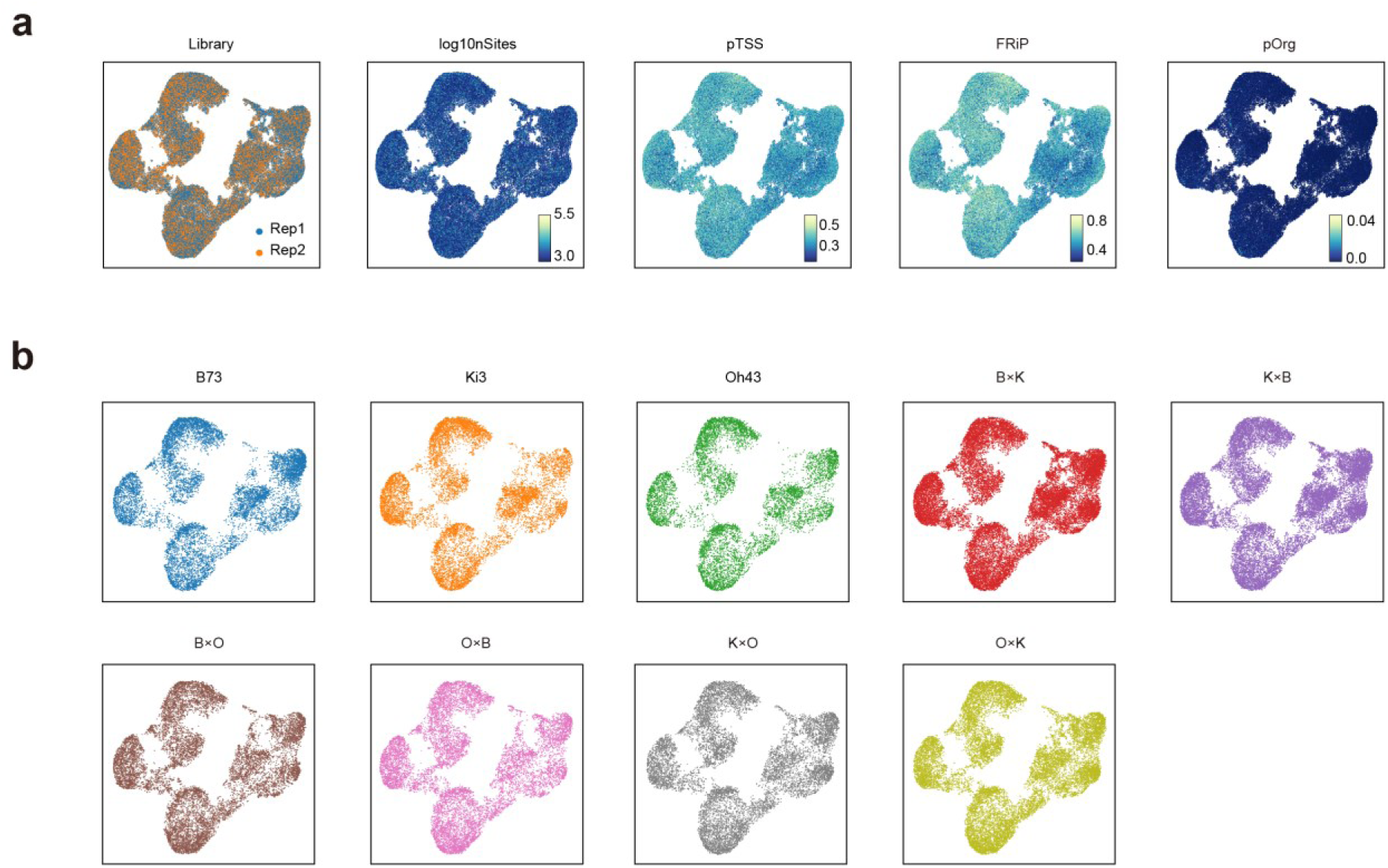
Assessment and quality control of scifi-ATAC-seq data for nuclei profiling in UMAP embeddings. **a**, UMAP visualization of scifi-ATAC-seq data showing, from left to right, library replicates, log_10_-transformed number of accessible sites (log_10_nSites), proportion of reads at TSS (pTSS), fraction of reads in peaks (FRiP), and proportion of organellar reads (pOrg). **b**, UMAP of all nuclei across the nine genotypes.

**Supplementary Fig. 5.**
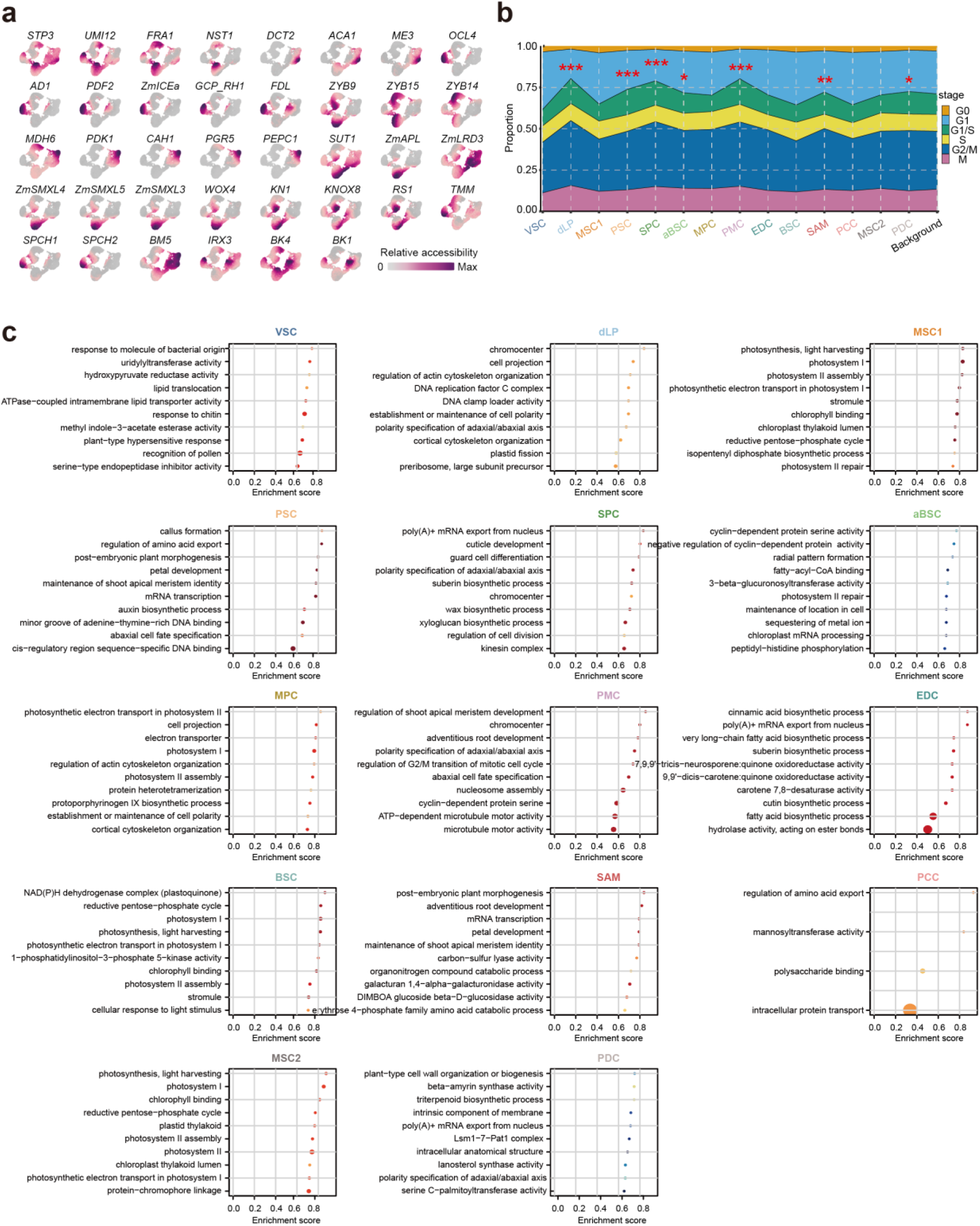
Cluster annotation. **a**, UMAP of nuclei colored by accessibility at cell-type marker genes, quantified across gene bodies and 500-bp upstream regions (gray, low; dark purple, high). **b**, Cell-cycle-stage proportions across cell types. Asterisks indicate enrichment relative to all 131,890 nuclei by hypergeometric test (* for *P* < 0.1, ** for *P* < 0.05, and *** for *P* < 0.01). **c**, GSEA bubble plots showing up to ten significantly enriched pathways per cell type, ranked by enrichment score. For aBSC and PDC, which had no significant pathways, the top ten pathways were shown. Dot color and size indicate adjusted P value and gene count, respectively.

**Supplementary Fig. 6.**
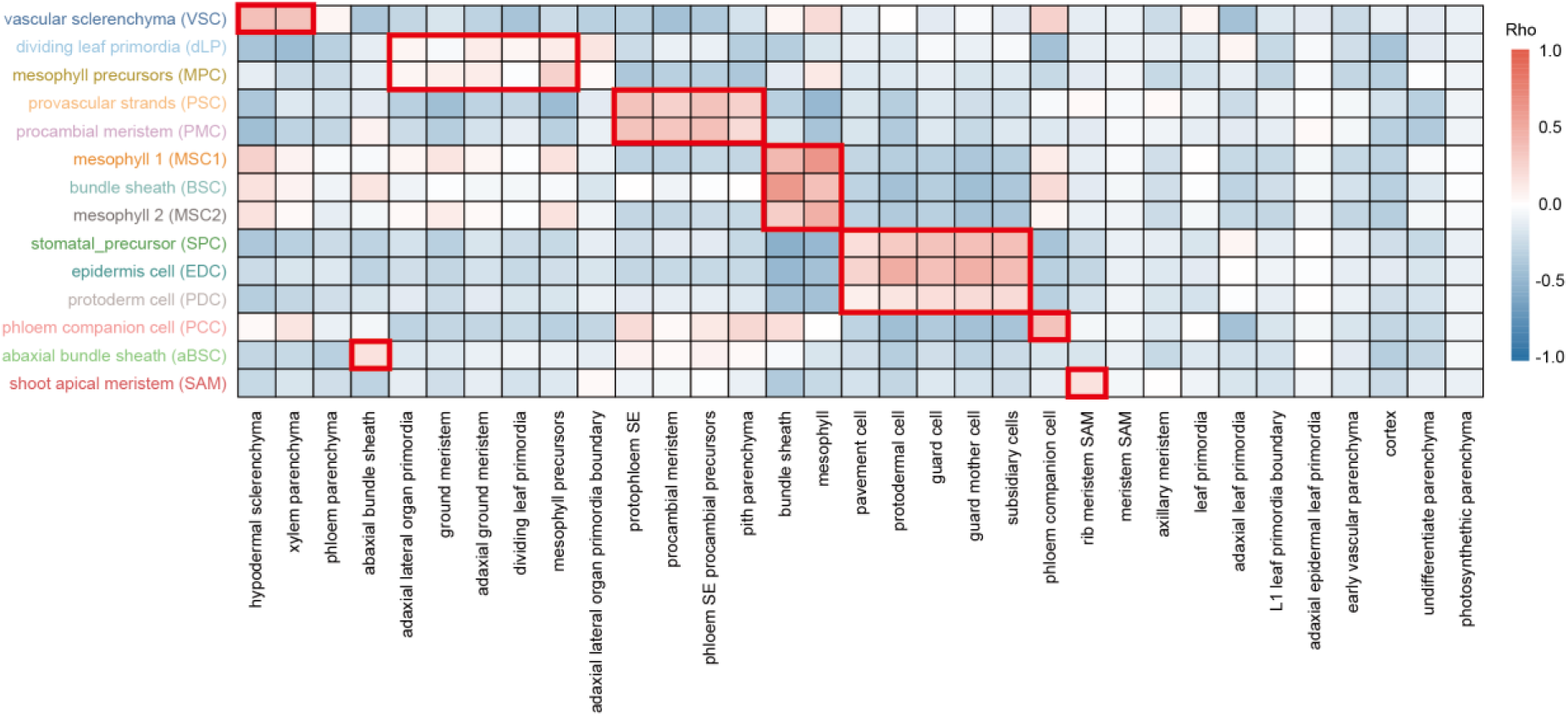
Cross-dataset comparison of chromatin accessibility profiles across cell types. Heatmap showing Spearman correlations between the 14 cell types identified in this study (rows) and previously annotated maize seedling cell types (columns) (*25*), based on row *Z*-scored CPM values across cell-type-specific ACRs (Tau > 0.7). Red boxes highlight the highly similar published cell type for each cell type identified in this study.

**Supplementary Fig. 7.**
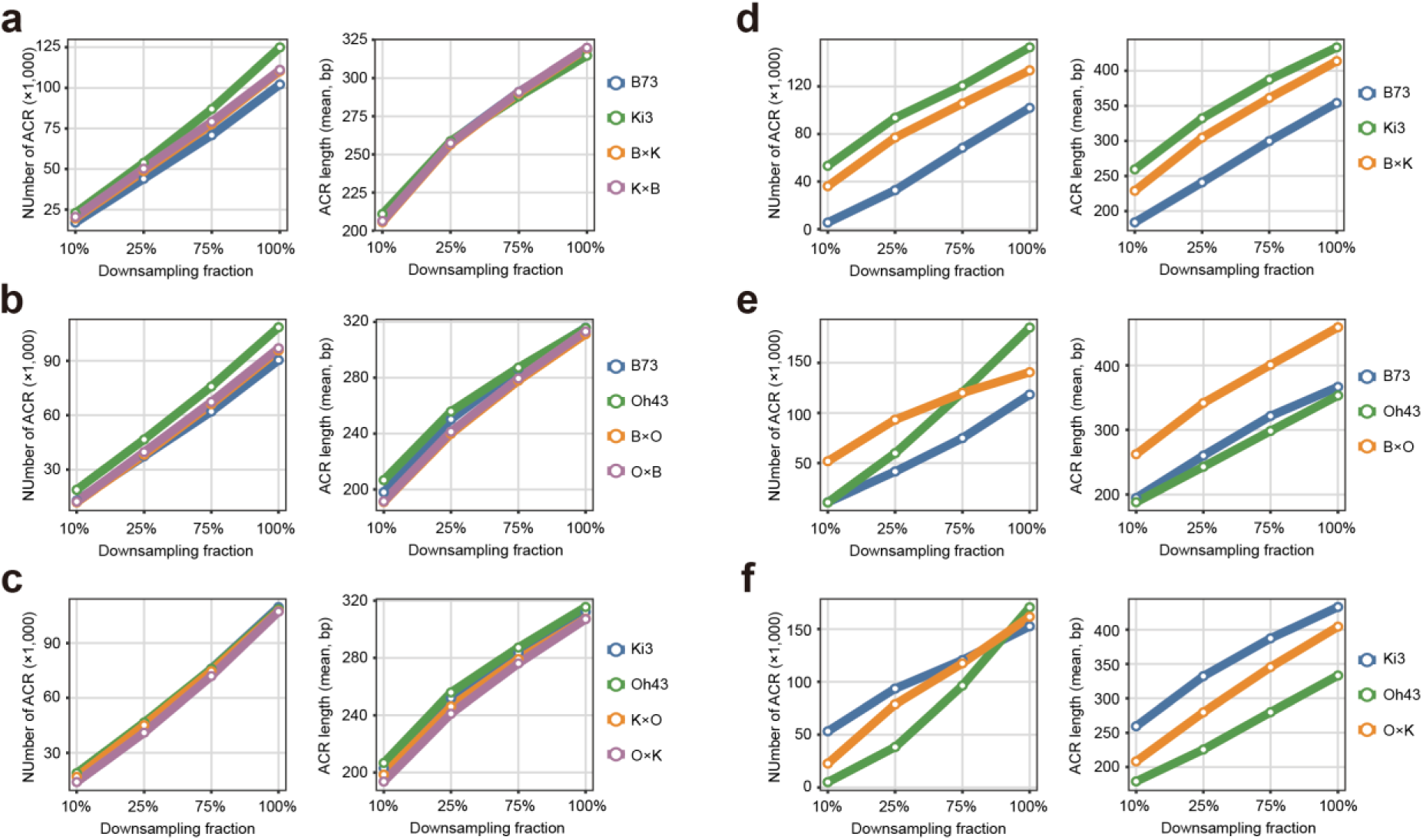
Matched-depth analysis of ACR number and length in inbreds and hybrids. **a–c**, Downsampling analysis of pseudo-bulk scifi-ATAC-seq data generated in this study for B73–Ki3 (**a**), B73–Oh43 (**b**) and Ki3–Oh43 (**c**) genotype comparisons. **d–f**, Re-analysis of published bulk ATAC-seq data (*8*) for the corresponding genotype comparisons. Reads were aligned to the B73 AGPv5 genome, filtered to reduce reference-mapping bias, and downsampled to 10%, 25%, 50% and 100% of the shared minimum Tn5 insertion depth before ACR calling. Points represent the mean value from five independent downsampling replicates. For each panel, the left plot shows the number of called ACRs and the right plot shows mean ACR length.

**Supplementary Fig. 8.**
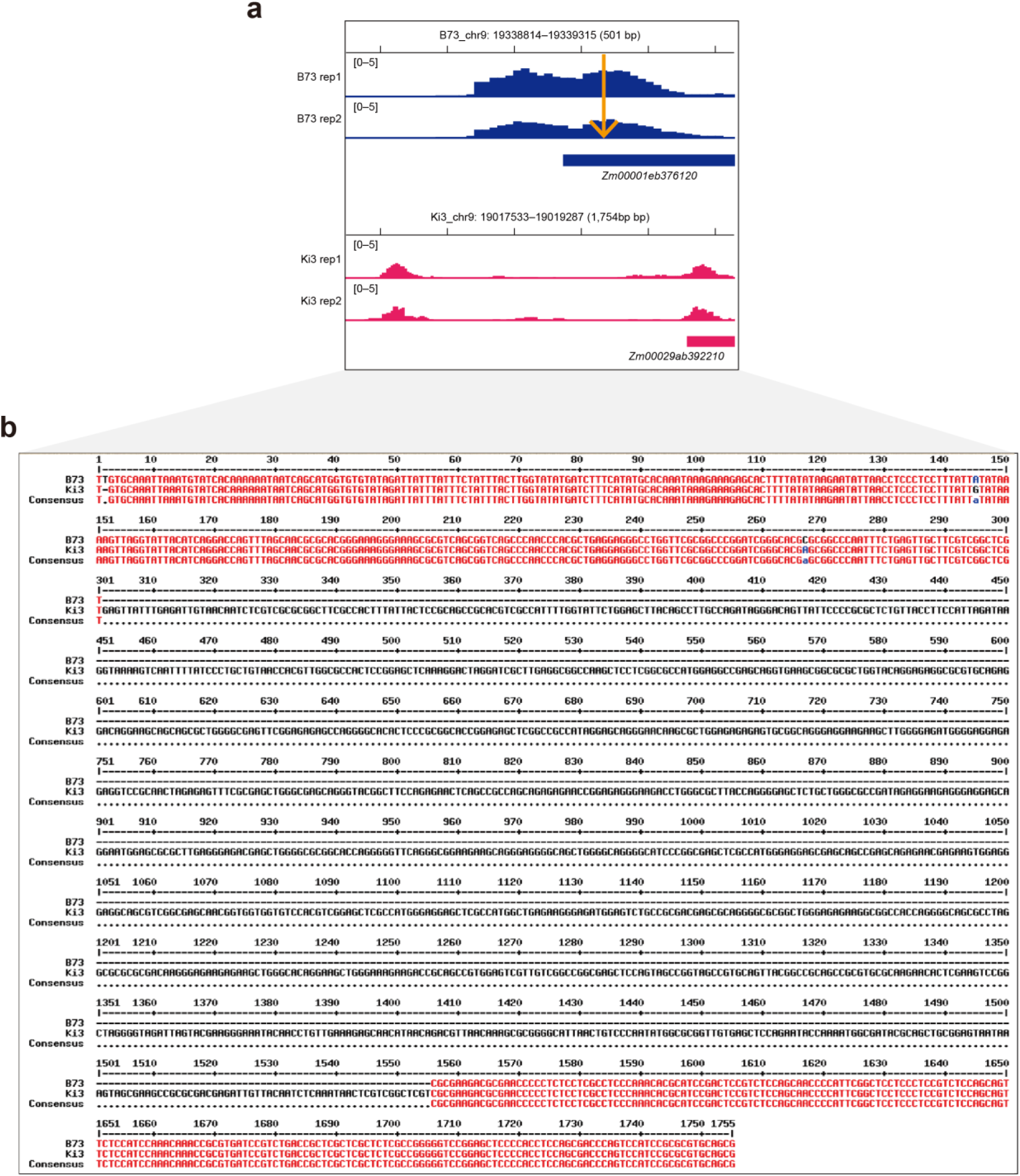
A representative example of an IS-ACR between maize inbred genomes. **a**, Genome browser views of an ACR locus showing chromatin accessibility profiles in B73 (blue) and Ki3 (magenta). Two biological replicates are shown for each genotype. While the overall accessibility pattern is conserved, the corresponding region in the Ki3 genome contains a ∼1.2-kb insertion relative to B73, resulting in a positional shift of the ACR. The insertion site is located near the ∼300 bp position and is indicated by the orange arrow. Gene models are shown below the tracks. **b**, Pairwise sequence alignment of the corresponding genomic regions between B73 and Ki3. The alignment confirms the presence of a ∼1.2-kb insertion in the Ki3 sequence that is absent in B73. Conserved bases are indicated in red color, and the inserted sequence is highlighted by alignment gaps.

**Supplementary Fig. 9.**
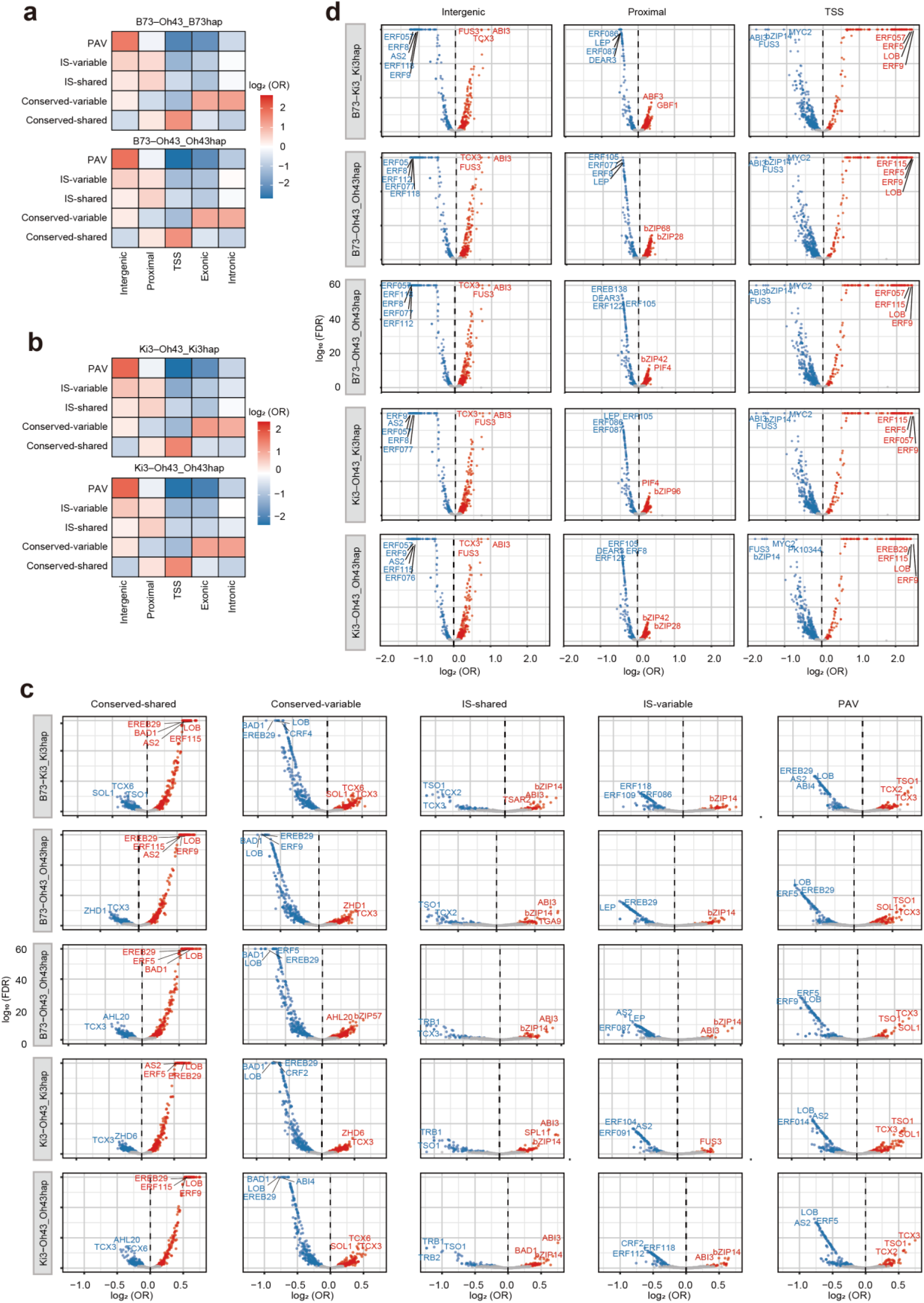
Genomic annotation and TF motif enrichment across ACR classes. **a–b**, Enrichment of ACR categories across genomic annotation classes in the B73–Oh43 (**a**) and Ki3–Oh43 (**b**) haplotypes. Heatmaps show log_2_ odds ratios, with red and blue indicating enrichment and depletion, respectively. **c–d**, TF motif enrichment across ACR categories defined by sequence conservation and accessibility (**c**; Conserved-shared, Conserved-variable, IS-shared, IS-variable and PAV) or genomic context (**d**; intergenic, proximal and TSS) in the remaining five haplotypes. Volcano plots show log_2_ odds ratios and –log_10_ FDR values for individual motifs. Significant enrichment and depletion are shown in red and blue, respectively, and non-significant associations in grey. –log_10_ FDR values are capped at 60, with consistent axis scales within each panel.

**Supplementary Fig. 10.**
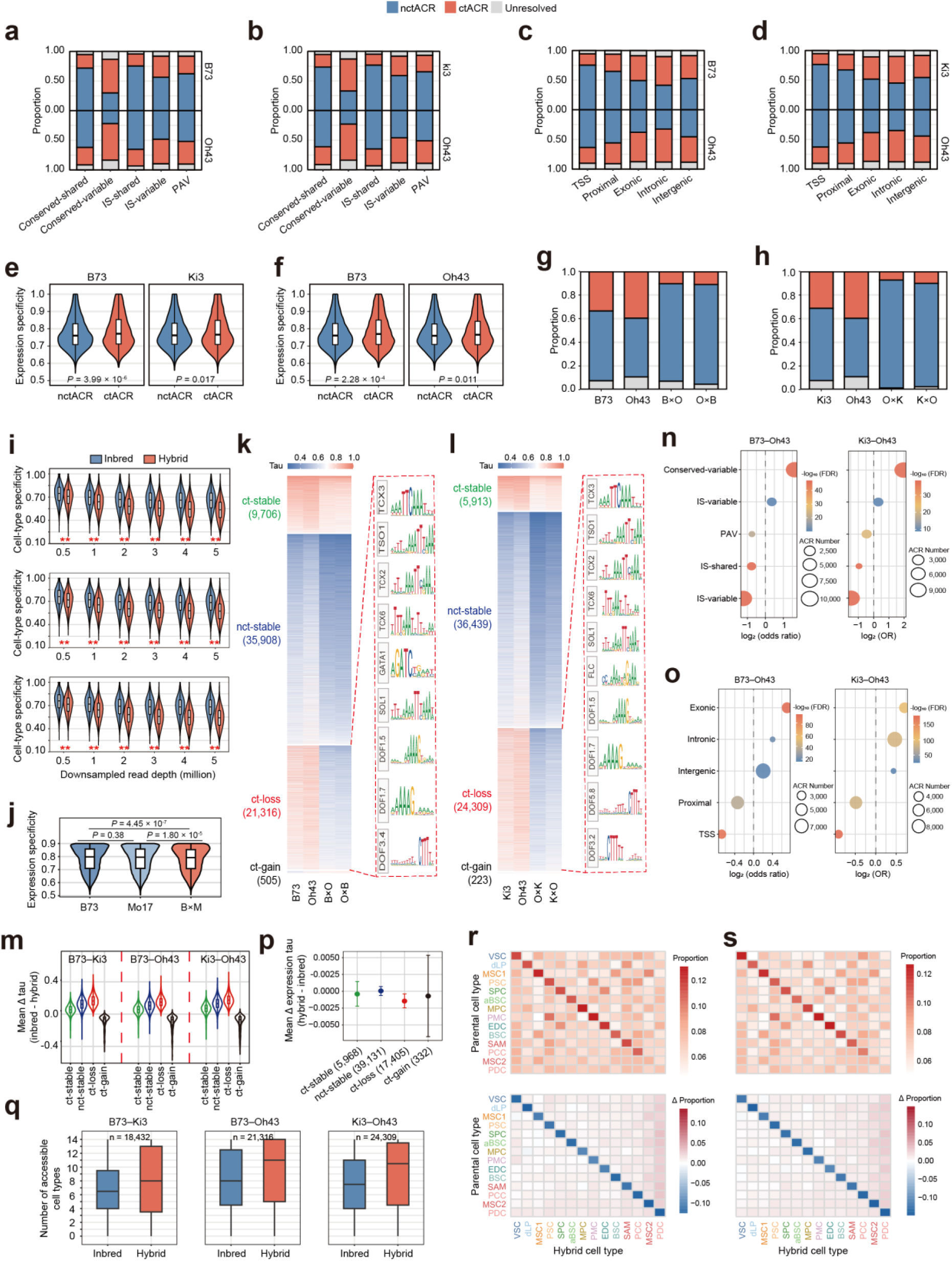
Hybridization attenuates cell-type-specific chromatin accessibility. **a–b**, ctACR, nctACR and unresolved ACR proportions across sequence/accessibility-defined ACR categories in the B73–Oh43 (**a**) and Ki3–Oh43 (**b**) comparisons, shown separately for each parental haplotype. **c–d**, ctACR, nctACR and unresolved ACR proportions across genomic contexts in the B73–Oh43 (**c**) and Ki3–Oh43 (**d**) comparisons. **e–f**, Expression specificity of genes linked to genic ctACR and ctACR for parental genotypes in the B73–Ki3 (**e**) and B73–Oh43 (**f**) comparisons. Expression specificity was calculated using previously published maize snRNA-seq data (*25*). **g–h**, ctACR, nctACR and unresolved ACR proportions in parents and reciprocal hybrids for B73–Oh43 (**g**) and Ki3–Oh43 (**h**). **i**, Cell-type specificity of ACRs after downsampling inbred and hybrid datasets to matched sequencing depths across the three parental–hybrid combinations. From top to bottom: B73–Ki3, B73–Oh43 and Ki3–Oh43. \*\**P* < 0.01. **j**, Tissue specificity of expression across 23 tissues in parental inbreds and their hybrid using an independent maize developmental transcriptome dataset (*47*). **k–l**, Tau heatmaps across parents and reciprocal hybrids showing four ACR specificity-state groups for the B73–Oh43 (**k**) and Ki3–Oh43 (**l**) comparisons, with representative enriched TF binding motifs shown on the right. **m**, Mean change in cell-type specificity (inbred minus hybrid) across four ACR specificity-state groups. **n**, Enrichment of sequence/accessibility-defined ACR categories in ct-loss ACRs for B73–Oh43 (left) and Ki3–Oh43 (right). **o**, Enrichment of genomic contexts in ct-loss ACRs for B73–Oh43 (left) and Ki3–Oh43 (right). **p**, Mean hybrid–inbred expression tau differences for genes nearest to ACRs in each specificity-state group. Points indicate group means; error bars indicate bootstrap 95% confidence intervals. **q**, Per-ACR accessibility breadth of ct-loss ACRs in inbreds and hybrids. Accessibility states were assigned using a two-component negative-binomial mixture model of replicate-level raw counts, with contexts having posterior probability ≥0.5 classified as accessible; breadth was defined as the number of accessible cell types and averaged across the two inbreds or reciprocal hybrids. n denotes the number of successfully fitted ct-loss ACRs included in each comparison. **r–s**, Hybrid accessibility profiles of ct-loss ACRs grouped by parental source cell type for B73– Oh43 (R) and Ki3–Oh43 (S). Top heatmap, mean accessibility proportion across hybrid cell types. Bottom heatmap, Δ proportion between hybrids and parents. Parental source cell type was defined as the cell type with the highest mean accessibility proportion across the two parents. Accessibility proportion was calculated for each ACR as the fraction of CPM-normalized accessibility assigned to each cell type across all 14 cell types and Δ proportion denotes hybrid mean minus parental mean proportion.

**Supplementary Fig. 11.**
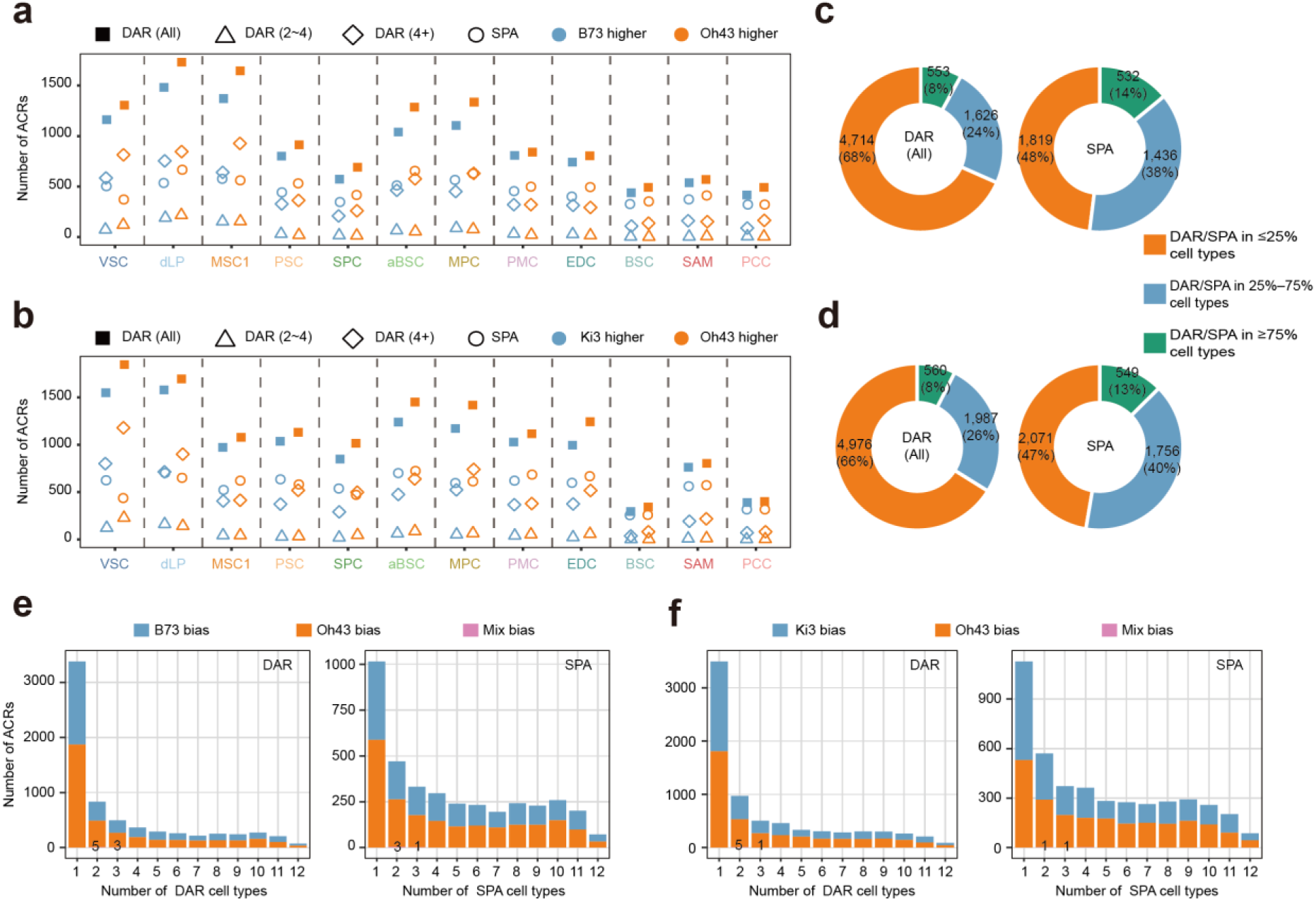
Cell-type-resolved parental divergence of chromatin accessibility between inbreds. **a–b**, The number of DARs in each cell type is shown, with parental bias indicated by color. DARs at increasing stringency (all DARs, 2-4, and 4+) are distinguished by different symbols. SPAs are shown by circles. Data are shown for the parental comparisons B73 versus Oh43 (**a**) and Ki3 versus Oh43 (**b**). **c–d**, DARs and SPAs were grouped by cell-type recurrence, defined as regions detected in ≤25%, 25%–75%, or ≥75% of tested cell types. Donut plots show the distribution of DARs (left) and SPAs (right) for the B73–Oh43 (**c**) and Ki3–Oh43 (**d**) comparisons. **e–f**, Numbers of DARs (left) and SPAs (right) detected across increasing numbers of cell types for the B73–Oh43 (**e**) and Ki3–Oh43 (**f**) comparisons. Colors indicate the directional consistency of parental accessibility divergence, with regions showing consistent bias toward one parent or mixed bias across cell types. Numbers above indicate ACRs with mixed bias and bars without labels contain no mixed-bias ACRs.

**Supplementary Fig. 12.**
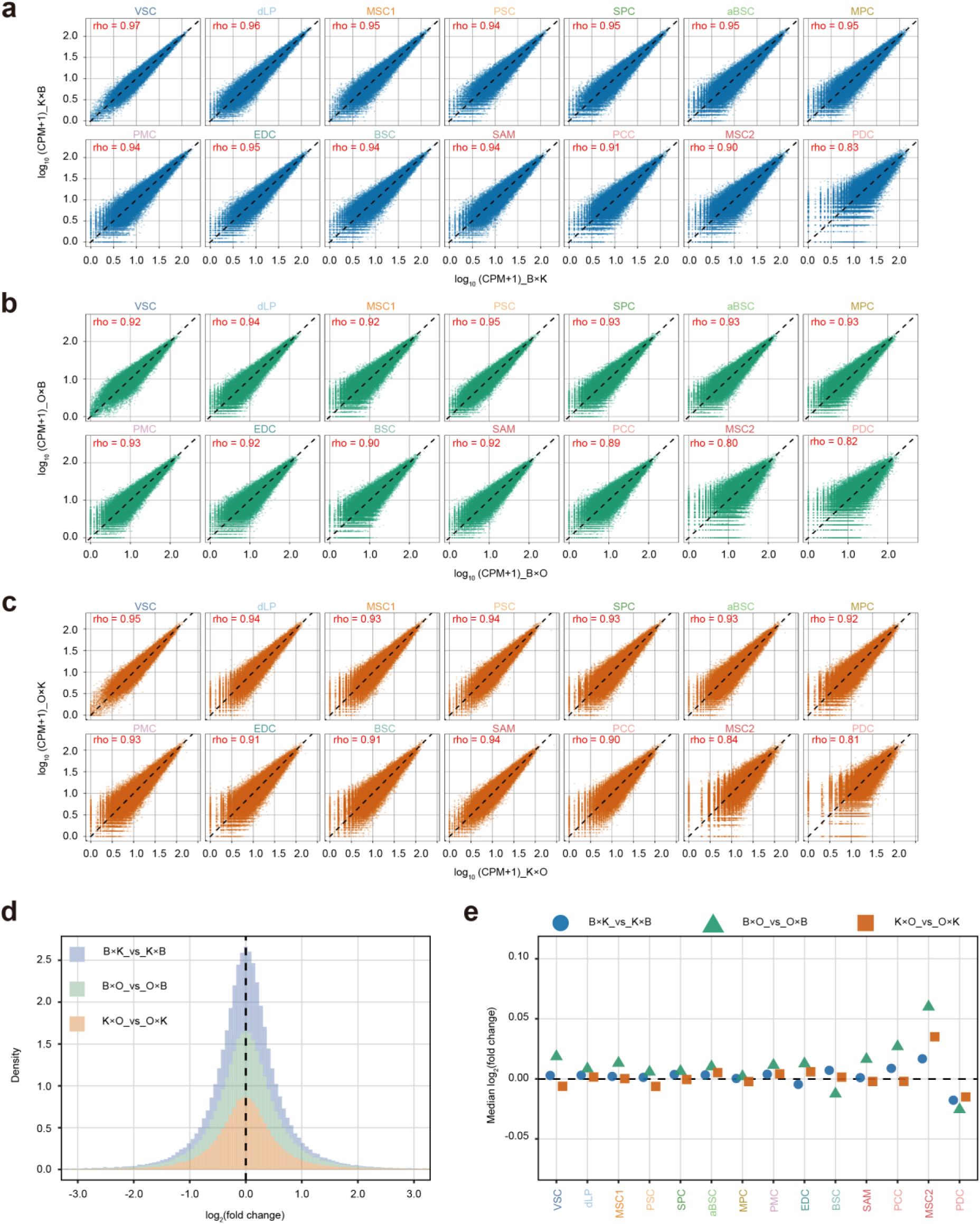
Reciprocal hybrids show no significant parent-of-origin effects. **a–c**, Chromatin accessibility in reciprocal hybrids across 14 cell types for B×K versus K×B (**a**), B×O versus O×B (**b**) and K×O versus O×K (**c**). Values represent replicate-averaged log_10_(CPM + 1), each point denotes an ACR, and Spearman correlation coefficients are shown. **d**, Distributions of log_2_ fold changes between reciprocal hybrids, pooled across cell types. **e**, Median log_2_ fold change across ACRs for each cell type and hybrid pair.

**Supplementary Fig. 13.**
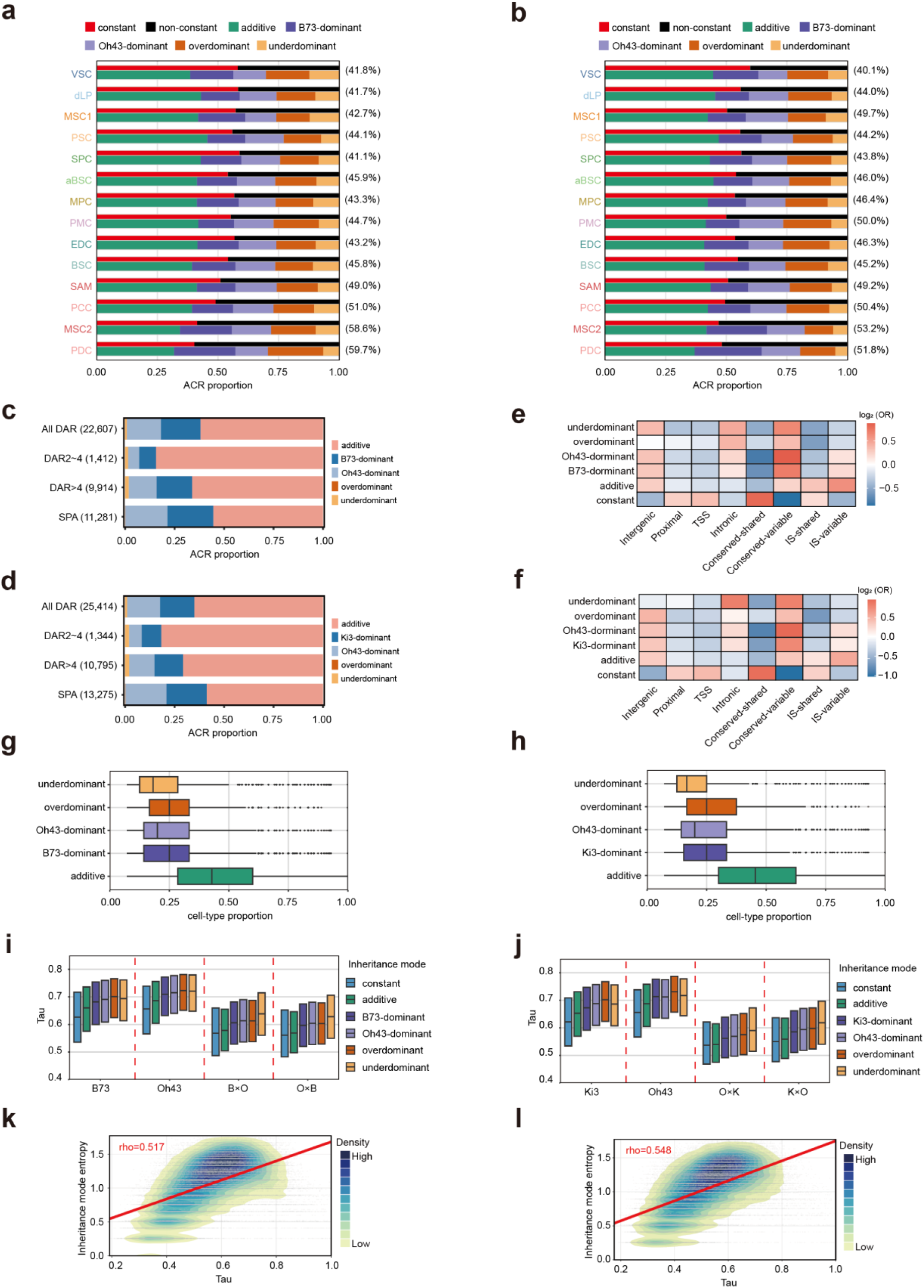
Inheritance-mode landscapes in B73–Oh43 and Ki3–Oh43. **a–b**, Cell-type-specific composition of ACR inheritance modes in the B73-Oh43 (**a**) and Ki3-Oh43 (**b**) hybrid comparisons. For each cell type, the upper bar shows the proportion of constant and non-constant ACRs, and the lower bar shows the composition of non-constant ACRs across additive, parental-dominant, overdominant and underdominant categories. Values in parentheses indicate the proportion of non-constant ACRs.**c–d**, Proportions of inheritance modes among ACRs in distinct parental DAR classes for B73–Oh43 (**c**) and Ki3–Oh43 (**d**). Values in parentheses indicate the total number of DARs pooled across cell types. **e–f**, Heat maps showing enrichment of inheritance modes across genomic contexts and ACR conservation/accessibility classes for B73–Oh43 (**e**) and Ki3–Oh43 (**f**). **g–h**, Breadth of non-constant inheritance modes across cell types for B73–Oh43 (**g**) and Ki3–Oh43 (**h**), quantified as the fraction of cell types in which each ACR–mode pair was observed. **i–j**, Tau distributions of ACRs grouped by inheritance mode across parents and reciprocal hybrids in the B73–Oh43 (**i**) and Ki3–Oh43 (**j**) comparisons. **k– l**, Density plots showing the association between mean hybrid Tau and Shannon entropy of inheritance modes across cell types for B73–Oh43 (**k**) and Ki3–Oh43 (**l**). Colors indicate local point density; Spearman’s rho is shown.

**Supplementary Fig. 14.**
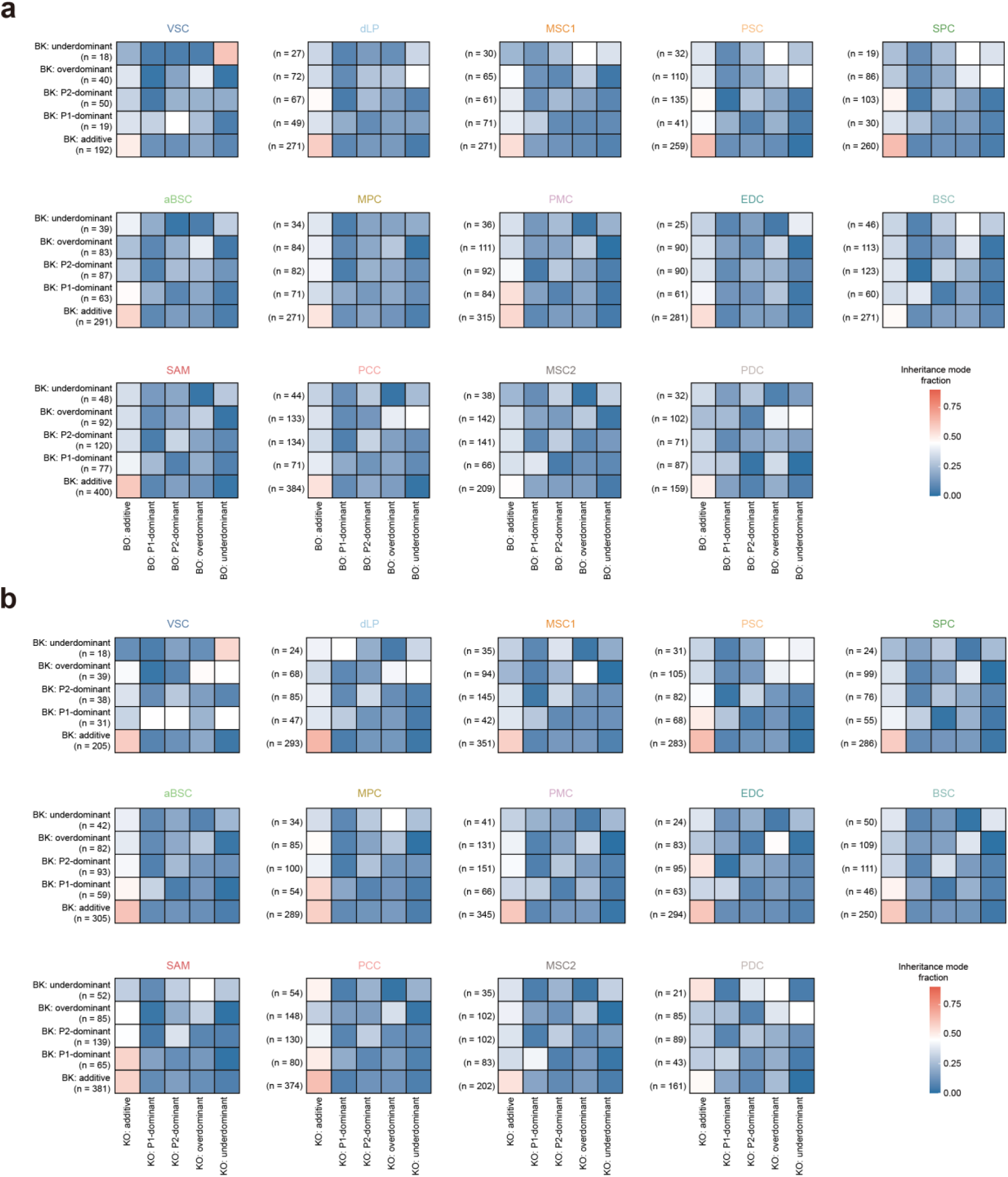
Cell-type-specific switching of inheritance modes across different hybrids. **a–b**, Transition rate matrices showing cell-type-specific changes of chromatin accessibility inheritance modes across hybrid combinations. Rows indicate inheritance modes in the reference hybrid and columns indicate the corresponding modes in the comparison hybrid. (**a**) BK vs BO. (**b**) BK vs KO. Colors denote transition rate. Numbers in parentheses indicate the number of loci assigned to each row category. In dominant categories, P1 denotes the parent shared between the two hybrids (e.g., B73 in BK vs BO), whereas P2 denotes the parent that differs between the hybrids (e.g., Ki3 in BK and Oh43 in BO).

**Supplementary Fig. 15.**
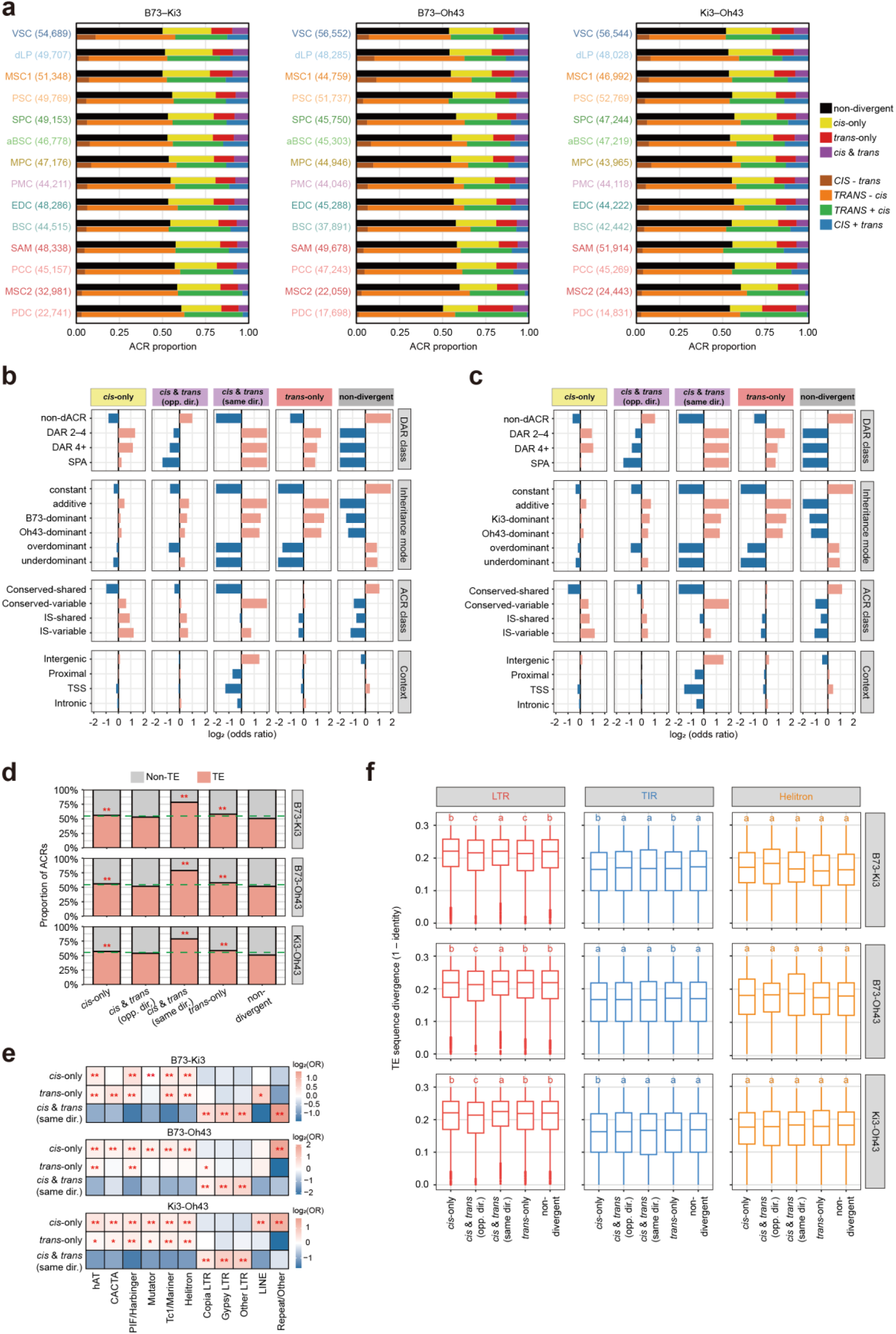
Overview of *cis*-and *trans*-regulatory pattern. **a**, Proportions of *cis*-and *trans*-regulatory categories across 14 cell types in B73–Ki3, B73–Oh43 and Ki3–Oh43 hybrids. Numbers in parentheses indicate classified ACRs per cell type. **b–c**, Enrichment of regulatory categories across parental DAR classes, inheritance modes, ACR sequence/accessibility classes and genomic contexts in B73–Oh43 (**b**) and Ki3–Oh43 (**c**). Bars show log_2_ odds ratios from Fisher’s exact tests; values beyond ±2 were capped at ±2. Red and blue indicate significant enrichment and depletion, respectively (FDR < 0.05), and grey indicates non-significant associations. **d**, Proportion of TE-overlapping ACRs within each regulatory category for each hybrid combination, restricted to intergenic ACRs. Dashed green lines indicate the overall TE-overlap fraction within intergenic ACRs. Asterisks denote significant enrichment relative to the remaining categories by Fisher’s exact tests. Only significant enrichments are marked: \*\**P* < 0.01. **e**, Enrichment of TE families among TE-overlapping intergenic ACRs in *cis*-only, *trans*-only and same-direction *cis* & *trans* categories. Colors show log_2_ odds ratios from Fisher’s exact tests. Only significant enrichments are marked: \**P* < 0.05; ** *P* < 0.01. **f**, TE sequence divergence (1 − identity) of LTR-, TIR-and Helitron-overlapping ACRs across regulatory categories. Boxplots are shown separately for each hybrid combination, letters indicate significant differences among categories.

**Supplementary Fig. 16.**
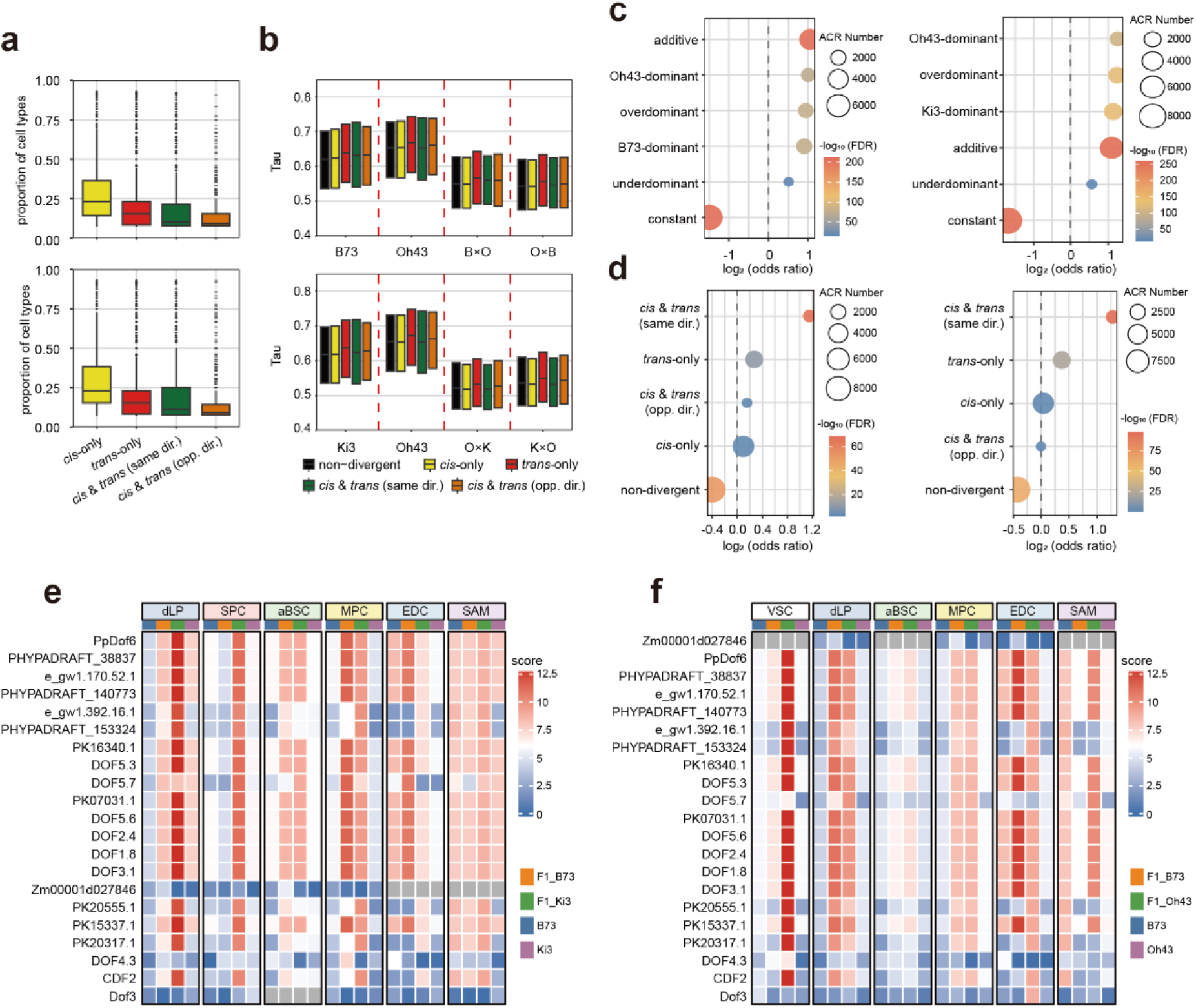
Cell-type-resolved regulatory features of *cis*-and *trans*-regulatory effects. **a**, Cell-type recurrence of regulatory effects in B73–Oh43 (top) and Ki3–Oh43 (bottom). For each ACR, recurrence was calculated as the fraction of cell types in which it was assigned to each category. **b**, Tau scores of ACRs assigned to each category across parental and hybrid genotypes in B73–Oh43 (top) and Ki3–Oh43 (bottom). **c**, Enrichment of ct-loss ACRs across inheritance modes in B73–Oh43 (left) and Ki3–Oh43 (right). **d**, Enrichment of ct-loss ACRs across *cis*/*trans* categories in B73–Oh43 (left) and Ki3–Oh43 (right). **e–f**, Heatmaps of normalized footprint scores for all DOF-family motifs surrounding the *GLK2*-associated ct-loss ACR (B73: chr3:1,590,027–1,590,528) in the B73–Ki3 (**e**) and B73–Oh43 (**f**) crosses. Only cell types with valid parent–hybrid footprint comparisons are shown. Motif names correspond to JASPAR motif IDs and grey color indicates missing measurements.

## Notes

### Competing Interest Statement

The authors have declared no competing interest.

